# Microtubule Probe for Correlative Super-Resolution Fluorescence and Electron Microscopy

**DOI:** 10.1101/351098

**Authors:** Xiaohe Tian, Cesare De Pace, Lorena Ruiz-Perez, Bo Chen, Rina Su, Mingzhu Zhang, Ruilong Zhang, Qiong Zhang, Qin Wang, Hongping Zhou, Jieying Wu, Pan Xiang, Bin Fang, Yupeng Tian, Zhongping Zhang, Giuseppe Battaglia

**Affiliations:** Huaxi MR Research Center (HMRRC), Department of Radiology, West China Hospital, Functional and molecular imaging Key Laboratory of Sichuan Province, National Engineering Research Center for Biomaterials, Sichuan University, Chengdu, China; School of Life Science, Anhui University, Hefei, P. R. China; Department of Chemistry, Anhui University, Hefei, P. R. China; Institute of Physical Science and Information Technology, Anhui University, Hefei, P. R. China; Department of Chemistry, University College London, London, United Kingdom; Institute for the Physics of Living Systems, University College London, London, United Kingdom; EPSRC/JEOL Centre for Liquid Phase Electron Microscopy, University College London, University College London; School of Materials Science and Engineering, Tongji University, Shanghai 201804, China; Biotechnology Centre, Anhui Agriculture University, Hefei 230036, China; CAS Center for Excellence in Nanoscience, Institute of Intelligent Machines, Chinese Academy of Science, Hefei, China; Institute for Bioengineering of Catalonia, The Barcelona Institute for Science and Technology (BIST), Barcelona, Spain; Catalan Institution for Research and Advanced Studies, Barcelona, Spain

## Abstract

We report a versatile cyclometalated Iridium (III) complex probe that achieves synchronous fluorescence-electron microscopy correlation to reveal microtubule ultrastructure in cells. The selective insertion of probe between repeated *α* and *β* units of microtubule triggers remarkable fluorescent enhancement, and high TEM contrast due to the presence of heavy Ir ions. The highly photostable probe allows live cell imaging of tubulin localization and motion during cell division with an resolution of 20 nm, and under TEM imaging reveals the *αβ* unit interspace of 45Å of microtubule in cells.

## Introduction

The structure of living cells is the result of a complex and highly dynamic assembly of proteins, lipids, and nucleic acids. Imaging techniques such as electron and optical microscopy have provided us with both structural and dynamic information necessary to dissect the cell complexity. Today we have fined tools such as x-ray crystallography and transmission electron microscopy (TEM) [1] to resolve the cell components structure down to atomic resolution. However, the new frontier of structural and cell biology is the understanding of cell units holistic combination and how their structure adapt to the very crowded and complex environment that is the cell interior [2, 3]. Techniques such as cryo-electron tomography [4] a provide us with a glimpse of how cellular components operate over multiple length-scales. New sample holders are allowing TEM imaging of wet samples and whole cells to be visualised[5]. Yet TEM lack of probes that enable molecular specificity and often suffers from a limited observation window. On the other hand, optical microscopy has been the workhorse of cell biology for more than a century as it can be combined with selective molecular probes such as fluorescent proteins, immunolabeling and synthetic dyes to image live specimens across several dimensions [6]. In the last two decades, super-resolution optical microscopy (SRM) techniques such as stimulated emission depletion (STED) and stochastic optical reconstruction microscopy (STORM) in particular are pushing the limits of optical microscopy resolution [7] down to tens of nanometres getting close to the TEM resolution. However SRM imaging is as good as the probe used and the ultimate resolution will always comprimise with the probe size. The most specific one, green fluorescent proteins and immune-labelling are at least few nm in size, DNA paint offer better versatility, but only organic dyes really could potentially push the limit below 1nm. In addition most SRM techniques require photostable, and for STORM photoswichable, fluorescent probes [6, 8], restricting the time resolution. All of these limit SRM to access the scale critical to reveal the molecular arrangements of cellular components [9] and thus bridging structure with function. A way around SROM and TEM limitations is to combine them to capture a multidimensional and correlated image that contains high resolution dynamic structural information alongside functionality, cellular localisation, and possibly chemical mapping [10, 11]. Correlative light electron microscopy (CLEM)[12, 13] is now becoming a critical tool to study biology and calls for a completely new way of designing probes that both possess the adequate photo-physical properties for SRM and yet with sufficient density to enhance contrast in electron microscopy [11]. Among the different chemistries, compounds containing at least one metal-to-carbon bond known collectively as organometallic molecules, are becoming extremely promising for CLEM [14]. In the past years we showed that metal complexes can be used for imaging in electron and optical microscopy nuclear DNA[15], mithochodrial DNA[16], lipid membranes[17] and nuclear factor kappa B [18]. Here we report the synthesis and characterization of a cyclometalated Iridium (III) complex (c-IrC) probe and its application for correlative STED and TEM imaging. We chose as subcellular target the microtubule, one of the major components of all eukaryotic cells cytoskeleton made by the polymerization of proteins known as tubulin. Such a protein comes in several isoforms with *α*- and *β*-dimers being the most common within the cell. Microtubule formation is catalysed by Guanosine-5’-triphosphate GTP binding[19, 20]. Microtubules act as scaffolds for most organelles as well as pathways for kinesin motors to facilitate active transport within cells. Microtubules regulate endosomes and lysosomes [21], axonal transport [22], DNA segregation and consequent cell division dynamics [23]. Tubulin role in mitosis has made it a critical target for an anticancer agent with several drugs designed to bind and inhibit microtubule formation, including nocodazole and taxols [24]. The design of probes with the ability to selectively target tubulin, without affecting its activity, is critical for the understanding its function and role in cellular processes. Iridium (III) complexes are ideal luminescent materials with tunable MLCT (metal-to-ligand charge transfer) emissive from blue to red, high photostability, and large stock-shift [25, 26, 27]. Taking advantage of these features, we designed the c-IrC probe with red emissions upon selectively binding tubulin in live cells, thus enabling high-resolution necessary for STED microscopy. Equally importantly, the high electron density of Ir atom allows for high contrast in TEM.

## Results and Discussion

We synthesized c-IrC complex (**Fig.1a**) and its two similar complexes (cIr-Tub and cIr-1, cIr-2) as control (**Fig.S1**) using vanillin and phenanthroline to coordinate the electron-dense Ir(III) and produce a complex with Ir-N and Ir-C bonds which possess high stability and photo resistance. Vanillin is a food additive with minimal cytotoxicity towards living systems that can form thiazole derivatives by carboxylation with o-aminothiophenol, resulting in larger conjugated systems and push-pull electronic groups. Phenanthroline is a classic coordination group with a planar conjugated system that enable yellow emission upon Iridium binding [27]. Such properties bestow the complex with longer lifetimes compared to more conventional organic fluoroscopes and, most relevant here, offer the capability for time-gated STED, resulting in improved imaging resolution[8]. Finally, thiazole derivatives and phenanthroline allow for hydrogen bonding and hydrophobic interactions (*π*– *π* and *π* -alkyl) with protein amino acid residues, which in turn restrain the molecular torsion of the extended conjugated systems augmenting the complex fluorescence and consequently enabling the so-called “light switch” phenomenon. The synthesis and characterization of the Ir (III) phenanthroline complex cIr-Tub and its analogous complexes cIr-1, and cIr-2 are provided in the supporting information. The final product was crystallized (**Fig.S2** and **TableS1**) and characterized by ^13^C NMR (**Fig.S2**)^1^ H NMR (**Fig.S3**) and MS spectroscopy (**Fig.S4**). The fluorescence spectrum in the aqueous condition (**Fig.S8)** showed that cIr-Tub exhibited a weak red broad emission from 520 nm to 700 nm [28], with a large stock-shift (118 nm) [29] and lifetimes in a 20 ns domain (**Table.S2**).

**Figure 1:**
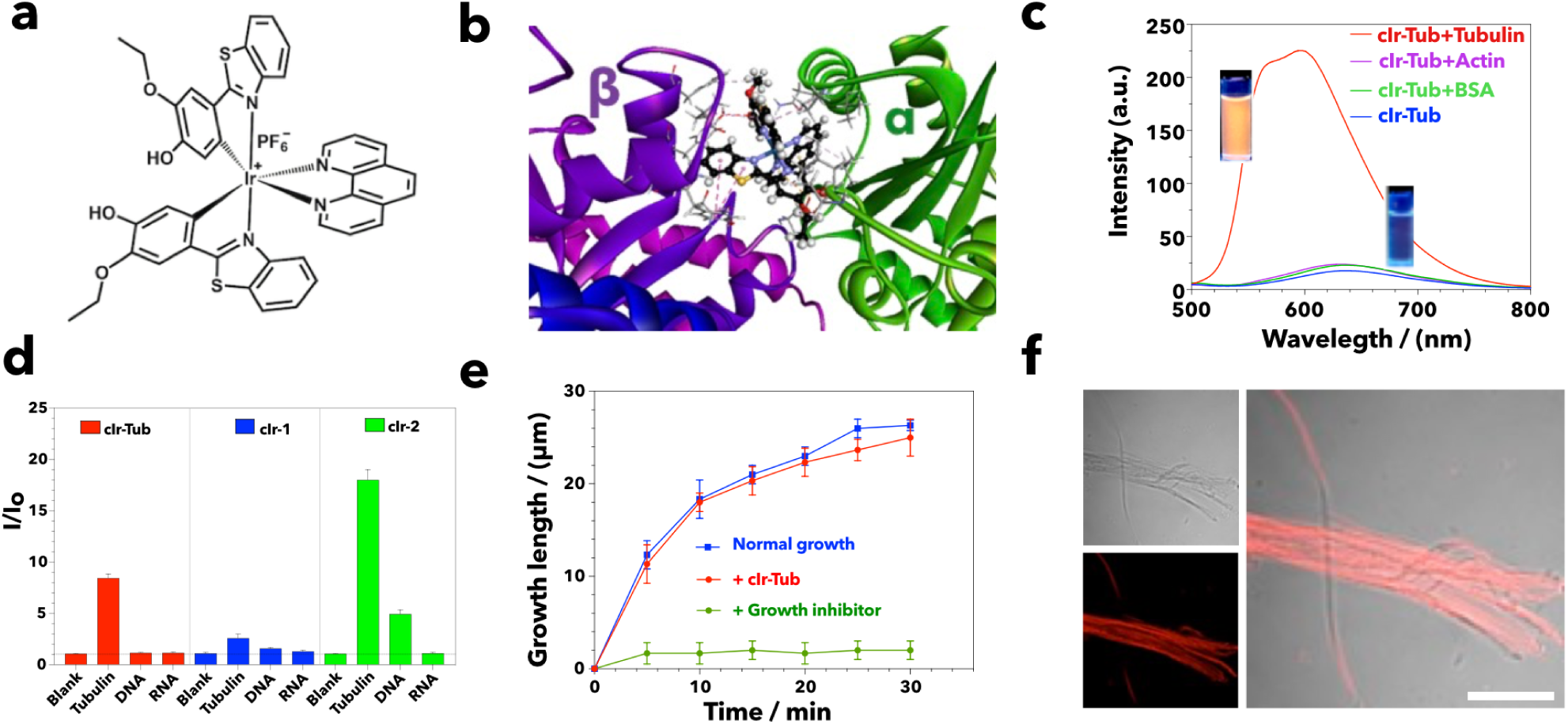
n vitro assessment of microtubules binding with cIr-Tub complex. . The chemical structure of cIr-Tub Ir complex (cIr-Tub) (**a**). Computer modelling of Ir-Tub complex (**b**) docking into between alpha and beta tubulin. Emission spectra of cIr-Tub complex (10 *μ*M) bound to tubulin, actin and bovine serum albumin (all concentration = 36 *μ*g/mL). The inserts show the corresponding cuvette of cIr-Tub and cIr-Tub + tubulin solution under UV irradiation (**c**). Normalised fluorescence intensity (**d**) of free (blank) cIr-Tub, cIr-1 and cIr-2 and the corresponding signal when bound to tubulin (30 μg/mL), DNA (10 μM), and RNA (10 μg/mL) respectively. In vitro polymerisation of tubulin protein into microtubules (**e**) in the presence of growth inhibitor (Nocodazole) and cIr-Tub with its correspondence florescent images (f), Scale bar = 20 *μ*m..

The binding of cIr-Tub to tubulin was first evaluated *in silico* via molecular docking using the Discover Studio ligand fit Vina software (version 2016, The Biovia Co) on the protein data base (PDB) tubulin heterodimer structure (5J2T) [30]. This particular PDB structure mimics the conformation of tubulin within microtubules in terms of structural rigidity yet the size is computationally manageable[30]. We set two tubulin dimers as the minimal units of receptor microtubules for cIr-Tub and show the resulting complex in **Fig.1b** as well as in **Fig.S6** and **Fig.S7**). We found that the Ir(III) complex binds to the tubulin dimer in the same site as vinblastine (**Table S3**), a well-known tubulin-targeting anticancer drugs [31]. As shown in **Fig.S6** cIr-Tub interacts with several tubulin residues including Phe214, Asn329, Pro325, Val353, Leu248, Val250, Tyr224, Leu227 and Tyr210 via *π*-*π* and *π*-alkyl interactions as well as hydrogen bonding mapped in **Fig.S7**. The interaction is relatively strong with binding energy circa 10 folds higher than that of vinblastine as reported in **Table.S3**.

We confirmed such interaction experimentally by fluorescence spectroscopy and observed that the cIrTub fluorescence is enhanced (**Fig.1c**) when the probe is bound to tubulin. As shown in our titration experiments in **Fig.S9** and polyacrylamide gel electrophoresis of tubulin protein (55 kDa) after staining with cIr-Tub (**Fig.S10**), both effects depend on the tubulin concentration. In **Table.S2** we report the measured photo-physical properties of the cIr-Tub fluorescence in different solvents and when bound to tubulin. The data show that the cIr-Tub bound to the protein has similar emission and excitation wavelengths as when dissolved in water but with lifetimes similar to those measured in dichloromethane. Most notably, while the cIr-Tub has an inferior quantum yield in aqueous solution when complexed to tubulin, its quantum yield is almost one order of magnitude higher than in any other solvent. Both the docking simulations and fluorescence spectroscopy suggest that binding extends to the overall *π*-conjugation system and inhibits the twisting movement of the complex. Such an interaction, in turn, hinders the non-radiative transition, resulting in augmented fluorescence which combined with the poor emission in water demonstrate a clear ‘light switch’ process ideal for probing the targeted proteins.

To confirm that both cIr-Tub binding and the light switch effect are selective to tubulin, we incubated the complex and the two control cIr-1 and cIr-2 complexes with different possible targets including bovine serum albumin (BSA), cytoskeleton protein actin, DNA and RNA. As shown in **Fig.1d** and **Fig.S11**, we observed a fluorescence increase only when cIr-Tub was incubated with tubulin with no detectable increase with other molecules. Complex clr-1 exhibited a weak response against tubulin while clr-2 showed a stronger fluorescence increase once bound to tubulin, but it also showed a lower yet considerable light-switch effect when binding to DNA displaying less specificity than cIr-Tub. Both data indicate that the vanillin moiety is critical for controlling the binding with tubulin, and when modified with an extra methyl group such as in cIr-Tub, it shows the desired selectivity. However, the deanol functionality in cIr-2 seems to increases the light-switch effect and presumably the binding affinity, the complex also interacts with DNA. Such a latter interaction is likely to be driven by the phenanthroline ligands as demonstrated with other metal complexes [32, 33]. To fully assess the suitability of cIr-Tub as a probe, we monitored tubulin polymerization *in vitro* 37°C. The results showed in **Fig.1e**report the tubulin growth into microtubules measured by optical microscopy using well-known inhibitor nocodazole (500nM) as a control [34]. After 30min, nocodazole prevented the tubulin polymerization, while cIr-Tub did not alter microtubule growth while still exhibiting the desired light-switch effect as shown in **Fig.1f**. Next, we investigated the application of cIr-Tub complex effect on live cells and measured possible cytotoxicity test incubating cIr-Tub complex with several types of cells including immortalized cancerous cells (A549, HeLa, MCF-7, HepG2 and HEK) as well as primary normal cells human dermal fibroblasts (HDF) and human embryo liver fibroblasts (HELF). As shown in (**Fig. S12**), we demonstrated quite a minimal effect of cIr-Tub on cell viability over 72h (i.e. three days) suggesting the probe is not affecting cell metabolic activity. Upon treatment of living cells with cIr-Tub, the intracellular microtubules emitted a strong red fluorescence signal under 450nm excitation wavelength. The *in situ* emission spectra was recorded under lambda model and fitted with the *in vitro* spectra in solution for a range of wavelengths (**Fig. S13**). Co-staining experiment were alos performed using the tubulin probes SiR-Tubulin (**Fig. 2a**) Furthermore, *α* -tubulin antibody (**Fig. 2b**), demonstrated the high degree of colocalization between cIr-Tub probe and microtubules in live and fixed cells, respectively. Tubulin selectivity was also demonstrated by the lack of colocalization with other subcellular components including mitochondria (**Fig. 2c**), actin ((**Fig. 2d**)), endoplasmic reticulum (**Fig. S14a**) and lysosomes (**Fig. S14b**). The aforementioned cIr-Tub analogous, cIr-1 and cIr-2, also displayed capability for reacting with tubulin protein and stained microtubules in live cells (**Fig.S15**). As showed in fluorescence spectroscopy, cIr-1 displayed a much weaker emission than clr-Tub, while cIr-2 showed unspecific nuclear targeting. Most interestingly cIr-Tub probe has proven to be very efficient for monitoring microtubule structure during mitosis (Fig. 2c). HepG2 cells were co-stained with nuclear dye, Nuc-Red, which allowed negligible signal overlap and highlighted the transformation from chromatin to chromosomes (prometaphase), indicating the progression of cell division. cIr-Tub displayed labelled microtubules polymerized through centriole (prophase) to mitotic spindles (metaphase). Structural and numeric aberrations (excess of centrosomes) can be commonly observed in human tumours [35]. Hence the ability of Ir-Tub to effectively monitor microtubules dynamically during mitosis provides an attractive feature for cancer biology investigations.

**Figure 2:**
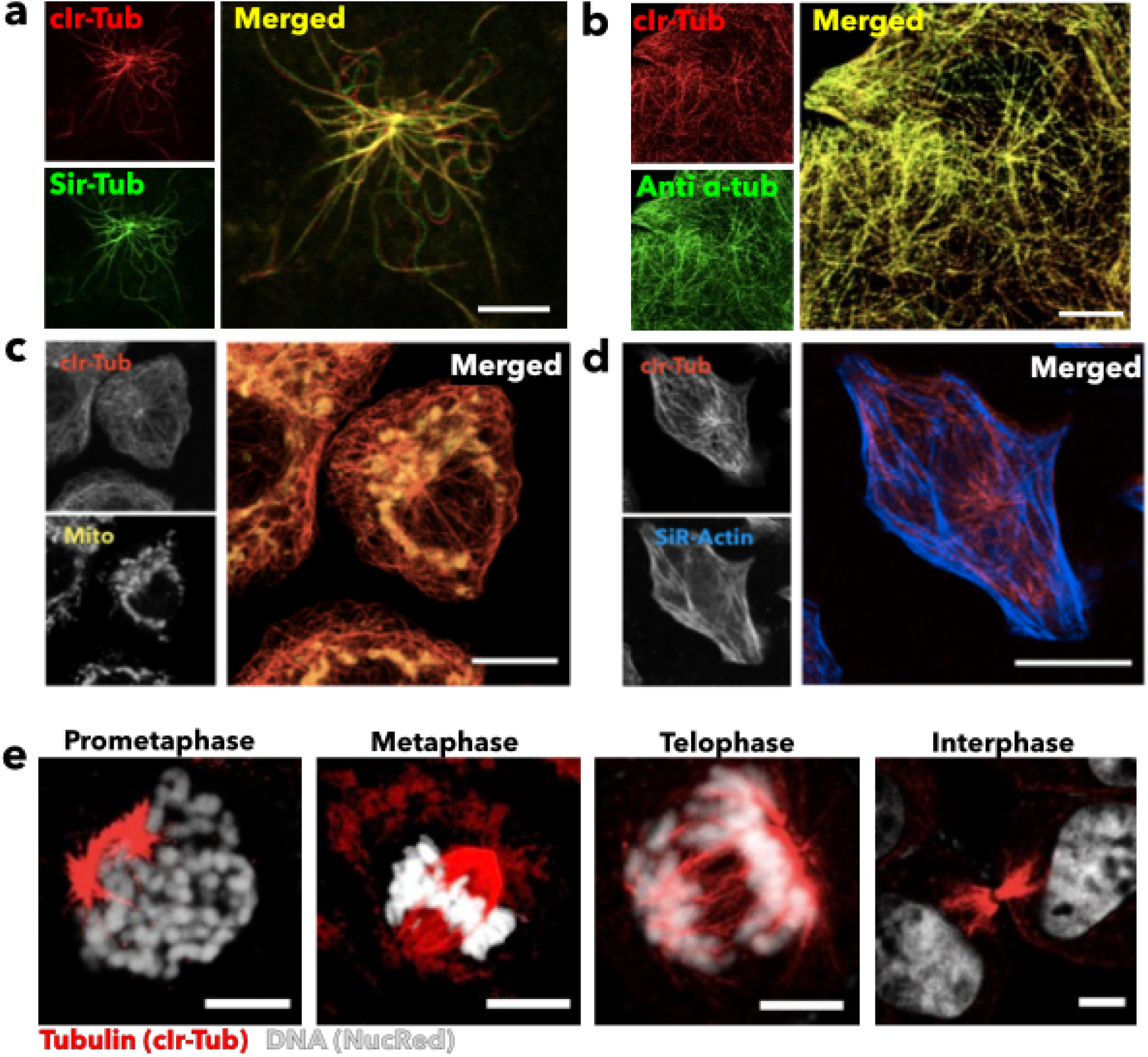
*In cella* assessment of cIr-Tub complex for microtubules staining. Confocal laser scanning micrographs of live HepG2 cells (**a**) incubated with 5*μ*g/mL cIr-Tub for 30 min and co-stained with SiR-Tubulin (*λ*_*Ex*_ = 640nm, *λ*_*Em*_ = 670nm). Confocal laser scanning micrographs of fixed HepG2 cells (**b**) incubated with 5μg/mL cIr-Tub for 30min and immunofluorescently labeled with *β*-Tubulin antibody (secondary: *λ*_*Ex*_ = 488nm, *λ*_*Em*_= 520-550nm) Scale bar=5*μ*m. Confocal microscopy images of live HepG2 cells treated with cIr-Tub and mitotracker (**c**), SiR-Actin (**d**) to mark the cell mitochondria and the other cytoskeleton protein, the actin, Scale bar=10*μ*m. Snapshots of dividing cells at different cell cycle phase (**e**) stained with Ir-Tub (5*μ*M) and with DNA dye NucRed (*λ*_*Ex*_ = 633nm, *λ*_*Em*_= 650-700nm). Scale bar=5*μ*m.

Encouraged by the above results, we further assessed the capabilities of cIr-Tub to mark microtubules *ex vivo*, and its utilization under stimulated emission depletion (STED) microscopy was also explored. The specificity of cIr-Tub for staining microtubules in multiple organs was initially proved using an immunofluorescent method (**Fig. S16**). Photo-bleaching experiments suggested that the fluorescence signal was preserved after 400 scans continued either in confocal or STED mode (**Fig. S17**), indicating a superior light resistance compared to other reported microtubule dyes (e.g. SiR-Tubulin, **Fig. S18**). We expected the cIr-Tub iridium metal core to be considerably less sensitive to the high laser power associated with STED imagining and hence enabling high-resolution imaging. In **Fig.3a** we compare conventional confocal laser scanning microscopy with both STED and time-gated STED imaging using the same sample of fixed HepG2 cell stained with 5μM cIr-Tub. The spatial resolution is enhanced in STED and even more so in time-gated STED with the micrographs showing microtubules as crisp fibril patterns with a superb signal-to-noise ratio. As shown in the graph in **Fig.3b**, the Ir-Tub labelled microtubules full width at half maximum (FWHM) goes down with increasing the STED laser power intensity or the gating time (ns). At full power and longest gating time we can image with resolution as low as 30nm and with 10% STED laser power and 20ns gated time we can image as low as 50nm. Such an augmented resolution is a critical factor when it comes to image sensitive samples such as the microtubules-rich brain tissues showed in **Fig.3c**. Here, STED imaging **Fig.3d** allows to highlight the elongated fibril structures throughout the hippocampus region and collect over 150 optical slides which in turn allows visualizing the microtubule networks in 3D **Fig.3e**. Such an imaging approach allows visualising complex neuronal microtubular network including growth cone, nascent neurites and spines with high spatial resolution as shown in **Fig.3f**.

**Figure 3:**
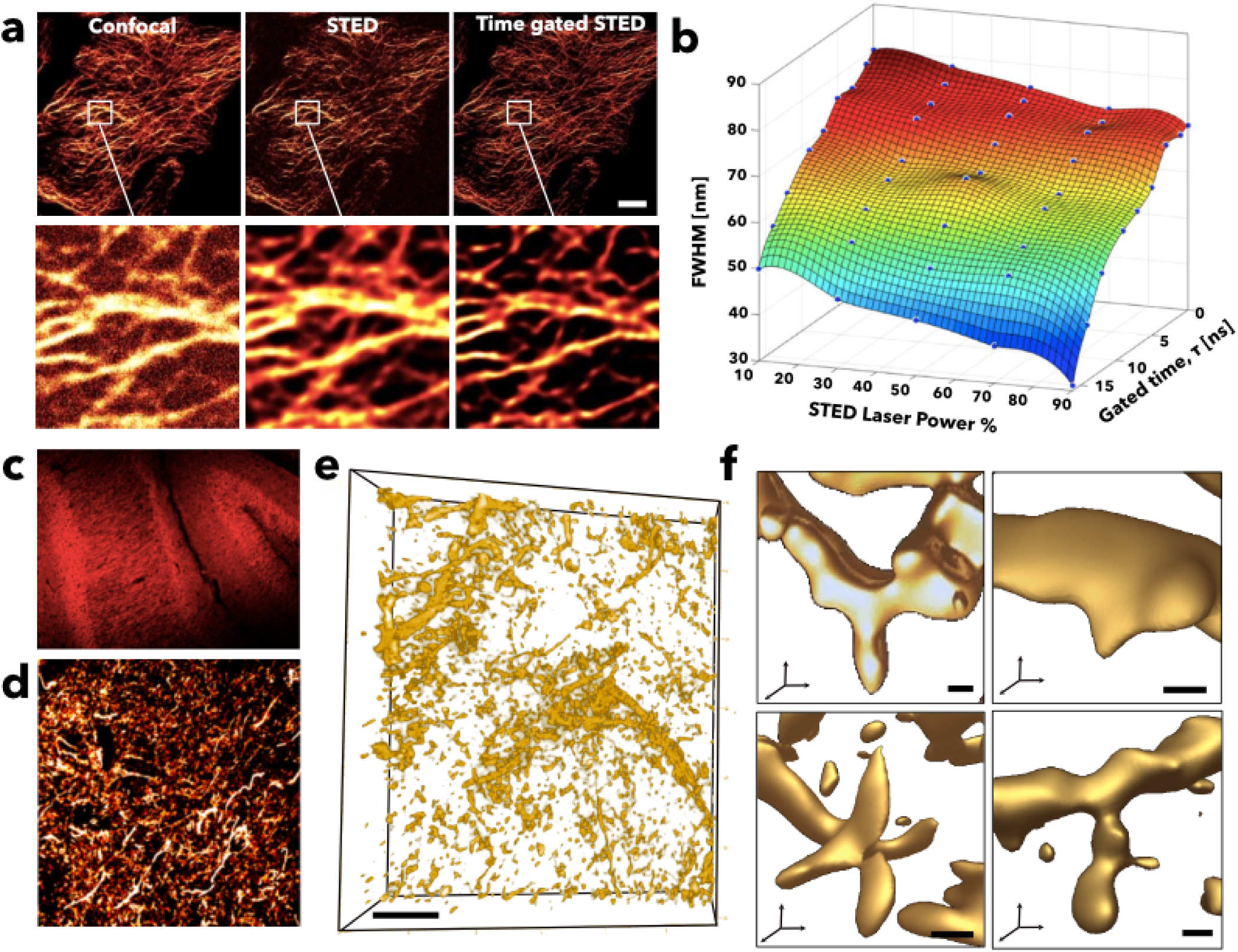
Feasibility of Ir-Tub complex for STED microscopy. HepG2 cell fixed and stained with 5μM Ir-Tub, sequentially imaged (**a**) by confocal, STED and time gated-STED (5ns) microscopy. Main micrograph scale bar = 10 *μ*m and region of interest scale bar =500nm. Graph (**b**) showing the Ir-Tub labeled microtubules resolution indicated by full width at half maximum (FWHM) as a function of the STED laser power intensity (donuts laser=592 nm, full power=12mW) and gating time (ns). Confocal micrographs of brain hippocampus sections labeled with cIr-Tub complex (**c**) and its magnified STED super resolution micrograph from neuronal rich region (**d**). Three-dimensional rendering of STED micrographs of hippocampus (**e**) and regions showing neuronal subunits from left to right: growth cone, nascent neurite, and spine ultrastructure (**f**). Scale bar=500nm.

STED microscopy is indeed increasing its capabilities for spatial resolution but, as shown in **Fig.3b**, it provides only tens of nanometres in resolution at best. While such a scale is sufficient to gather information about the morphology and connectivity of the neural network, as shown above, the actual microtubule molecular structure requires higher resolution not achievable yet by optical microscopy but well within the range of electron microscopy. Here the metal Iridium, the central part of the Ir-tub complex has sufficient high electron density to provide contrast [14] for TEM imaging. We thus incubated HepG2 cells with cIr-Tub complex and embedded them in epoxy resin, and microtome sectioned the resulting block into sections of 80nm thickness. We used conventional OsO_4_ as control stain and imaged the sections by transmission electron microscopy. As expected, the OsO_4_ treated cells displayed vesicles, endoplasmatic reticulum (ER) and mitochondria, however microtubule structure were not visible **Fig.4a**. Cells treated with cIr-Tub complex show almost uniquely **Fig.4b** and cells treated by both stains **Fig.4c** show a quite interesting complex network where the microtubules seem to interact intimately with organelles. Other stains like uranyl acetate (UA) or lead citrate (LC) can also stain microtubules, but they do this along with many cytosolic proteins and hence they lack of specificity [36]. The comparison between cIr-tub and UA/LC stain is shown in **Fig.S20** where we demonstrate that the image contrast by cIr-Tub not only provided a better resolution of the microtubules, but also a contrast enhancement compared to UA/LC stain.

**Figure 4:**
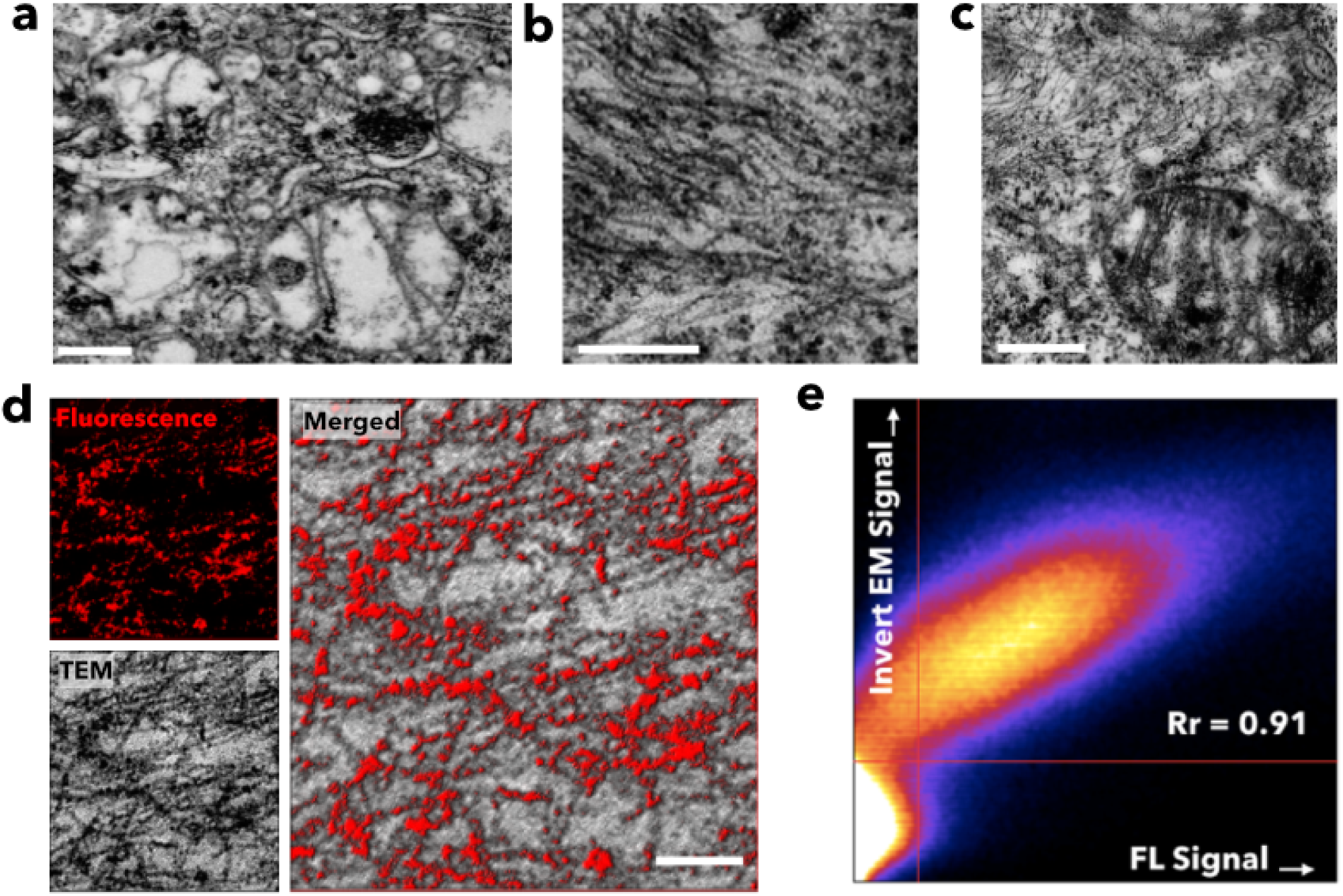
Correlative light and electron microscopy. Transmission electron microscopy micrographs of HepG2 cells treated with osmium tetroxide OsO_4_ (**a**), cIr-Tub (**b**) and OsO_4_+cIr-Tub (**c**), respectively. Scale bar = 500nm. Correlative STED fluorescent imaging and scanning-transmission electron microscopic image (**d**) of cIr-Tub labelled microtubules in HepG2 cells. The images obtained using light and EM microscopy were taken from the same site and the co localization scatter plot between fluorescent and inverted electronic microscopic signal (**e**) shows high correlation with Pearson’s coefficient, Rr =0.91.

Nevertheless, the dual metallic and fluorescent nature of cIr-tub allows an unprecedented combination of imaging modalities providing selective contrast for both fluorescence and transmission electron microscopy. We thus imaged cell sections using correlated light electron microscopy CLEM by sequential confocal microscopy (airyscan model) followed by scanning-transmission electron microscopy (STEM). As shown in **Fig.4d** the microtubule network was both visible under STEM and confocal imaging with the two signals overlapping with a colocalization Pearson coefficient of Rr=0.91 **Fig.4e**, confirming cIr-Tub affinity towards tubulin and showing its suitability for CLEM imaging. To further prove that the cIr-Tub is associated with the microtubules, cell sections were also imaged via energy-filtered transmission electron microscopy (EF-TEM). Such imaging mode allows chemical mapping and hence the identification of the Iridium atoms position within the microtubule structures by selecting electrons that have lost a specific amount of energy from inelastic scattering interacting with the inner-shell ionization. In the present study, the iridium O shell maps were acquired alongside conventional TEM imaging. In **Fig.5a** the detail of a single microtubule structure is shown both in conventional (unfiltered) TEM, in EF-TEM and a merged image showing the Ir elemental distribution along with the microtubule structures with subnanometer resolution. The distances between neighbouring Iridium atoms obtained from the elemental map were measured and plotted in **Fig.5b** showing that the probes were distributed along the microtubule structure with average interspaces of 4.2 nm. Such a value matches exceptionally well the distance estimated by placing the cIr-tub within the predicted binding socket as shown in the microtubule model reconstructed in **Fig.5c**. These results strongly suggest that Ir-Tub complex could be effectively internalized within intracellular microtubules and indeed was capable of acting as a multi-function probe for light (Confocal, STED, time-gated STED) and electron microscopy (TEM, STEM and EFTEM).

**Figure 5:**
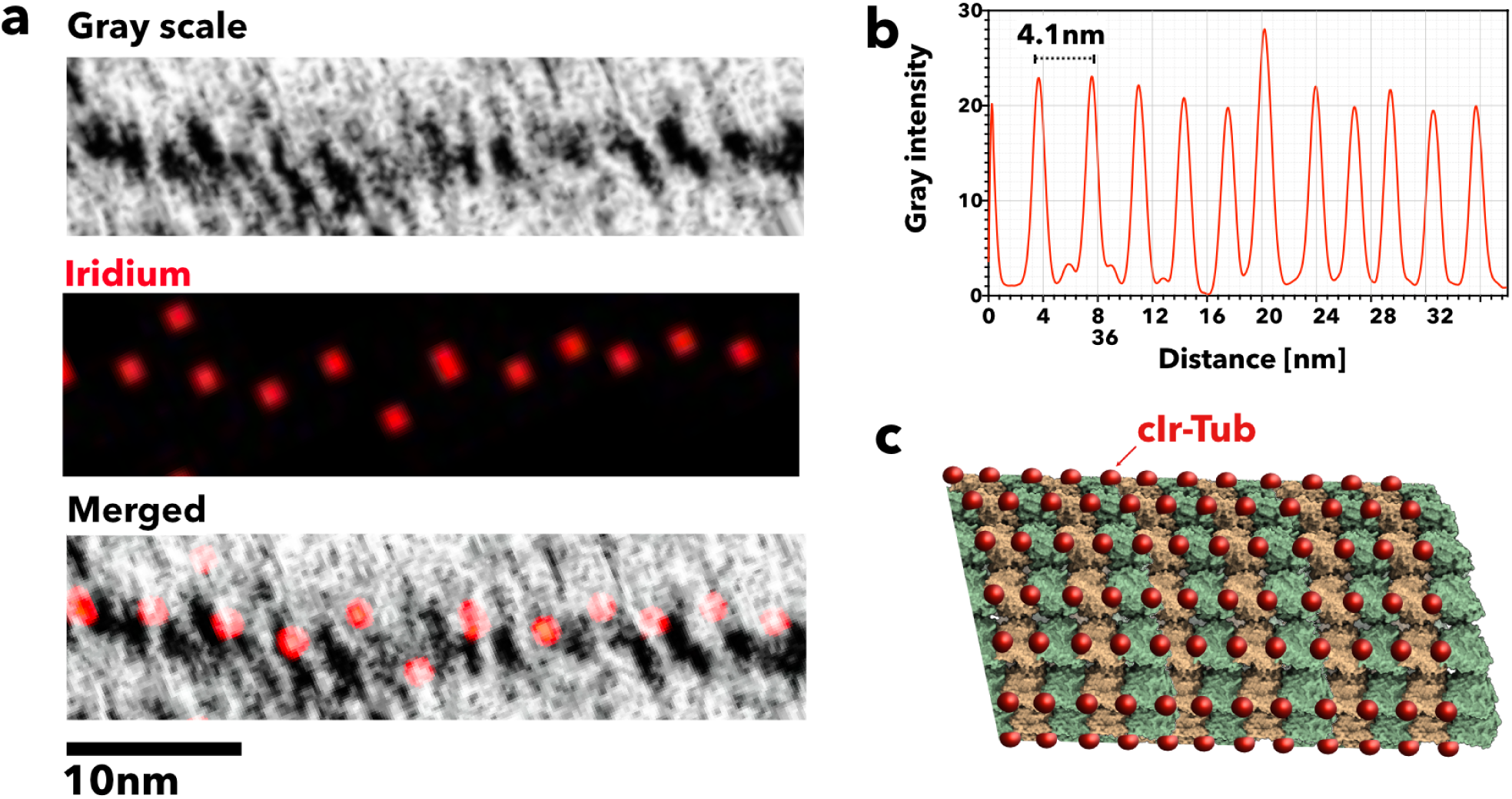
Energy filtered transmission electron microscopy (EF-TEM) imaging. Conventional TEM image, Iridium elemental mapping and merged images (**a**) of a single intracellular microtubule. The spacing between the Ir signals in the microtubule is periodic (**b**) with an average period length of 4.1 nm. This measure matches the distance between the Ir-Tub complex (red sphere) tubulin binding sites as shown in the schemes (**c**) of the microtubule reconstructed from the protein database structure 3J2U.

## Conclusions

In summary, we have designed and synthesized an Iridium (III) complex, cIr-Tub, that is capable of binding to *αβ*-tubulin active pockets. The MLCT emission of the cIr-Tub complex ‘switched on’ upon tubulin-binding allowing high specificity and sensitivity *in vitro* studies. *In cella* investigations firstly demonstrated that the proposed probe not only enables microtubules tagging and monitoring of several cell phases during mitosis but also shows minimal interference for cell proliferation over days. In particular, it was demonstrated that microtubules could be observed using cIr-Tub at *ex vivo* tissue level, and further STED micrographs displayed excellent signal-to-noise ratio. Ultimately, cIr-Tub complex offered unique, specific electron contrast for several TEM techniques (TEM, CLEM, EF-TEM). The iridium complex presented herein offers unique properties to be used as a fundamental tool to further understand microtubule dynamics in living systems, as well as a potential targeting vector for therapeutic purposes.

## Acknowledgements

X.T Y.T. and Z.Z. thanks the National Natural Science Foundation of China (51432001, 51672002, 21871003 and 51772002) for funding most of the experimental work. GB thanks to the EPSRC (EP/N026322/1) the ERC (CheSSTaG 769798) for funding part of his salary. C.D.P and L.R.P. thank Jeol for sponsoring their salary and sponsoring part of this work. XT and GB thank that Anhui 100 Talent program for enabling people travelling and sample exchange across the two sites.

## Author contributions

X. T., GB, Z.Z and Y.T design the project. R. Z., Q. Z and Y.T design and synthesized the complex. R. Z., Q. Z and ZZ. did all the characterization and data analysis. XT, R. S., and MZ. performed the cell and tissue florescent work, including confocal and STED. QW performed TEM sample preparation and sectioning. BC performed the CLEM experiment and related data analysis. C. D. P., L.R.P. and GB performed the EF-TEM and analysis the data. XT, Z.Z, CDP, L.R.P. and GB wrote the paper.

## Supporting information

### Materials

Dimethyl sulfoxide (DMSO), N, N-dimethylformamide (DMF), ethanol, methanol, dichloromethane (DCM) and NH4PF6 were purchased from Shanghai Chemical Reagents Co. Ltd (China). Ethylvanillin, 2-aminobenzenethiol, phenanthroline and Iridium chloride trihydrate were supplied from Aladdin Co. Ltd (China). Adenosine triphosphate (ATP), 5-dimethylthiazol-2-yl-2,5-diphenyltetrazolium bromide (MTT) were obtained from Beyotime Biotech. Co. Ltd. (China). Cell staining Dyes, 4’,6-diamidino-2-phenylindole (DAPI), Lysotracker Red, Mitotracker Deep-Red and endoplasmic reticulum tracker (ER tracker blue) were commercial products from Thermo Fisher Scientific (USA). SiR-Actin (Far-red) and SiR-Tubulin (Far-red) for live-cell cytoskeleton labelling were provided by Cytoskeleton, Inc. (USA). Anti-alpha Tubulin antibody and its secondary antibody (Alexa-488) were purchased from Abcam (China).

### Instruments

IR spectra were recorded on a Nicolet FT-IR 170 SX spectrophotometer (USA). The 1H-NMR and 13C-NMR spectra were obtained on Bruker AV 400 spectrometer (Germany) with TMS as an internal standard. Mass spectra were performed on a Micromass GCT-MS spectrometer (UK). UV-vis absorption spectra were recorded on a SHIMADZU UV-3600 spectrophotometer (Japan). Fluorescence spectra were collected with a HITACHI F-7000 spectro-fluorophotometer (Japan). Mouse tissue was sectioned under Leica CM3050 S system (Germany). Fluorescent microscopy was performed on a Leica SP8 (Germany) equipped with White Laser. STED nanoscopy experiments were performed under Leica DMi8 confocal microscopy equipped with Leica TCS SP8 STED-ONE unit. Transmission electron microscopy was carried out on a Hitachi HT7700 120kV Compact-Digital TEM (Japan). Energy-Filtered TEM for chemical analysis was done on JEOL JEM-2200FS TEM (Japan) equipped with a field emission gun (FEG) at 200 kV, and an in-column energy Omega filter. Correlative Light and Electron Microscopy (CLEM) experiments were operated using ZEISS LSM800 with Airyscan (Germany) and GeminiSEM 500 scanning transmission electron microscopy (Germany), correlated by ZEISS ZEN blue software.

### X-ray crystallography

The X-ray diffraction measurements were performed on a Bruker SMART CCD area detector using graphite monochromated Mo-Kα radiation (*λ* = 0.71069 Å) at 298(2) K. Intensity data were collected in the variable *ω*-scan mode. Direct methods and difference Fourier transformations solved the structures. The non-hydrogen atoms were refined anisotropically, and hydrogen atoms were introduced geometrically. Calculations were performed with the SHELXTL-97 program package.

### The synthesis of cyclometalated Iridium (III) complexes

cIr-Tub and its analogous cIr-1 and cIr-2, are detailed in **??**. Synthesis of ligand 4-(benzo[d]thiazol-2-yl)-2-ethoxyphenol. A mixture of 3-ethoxy-4-hydroxy benzaldehyde (3.32g, 20 mmol) and 4-aminothiophenol (2.50 g, 20 mmol) were dissolved in 50 mL of ethanol, and heated to reflux for 6 h. After cooled down to room temperature, the yellowish crystal powder was obtained by filter (4.91 g, yield 90.5 %). 1H NMR (400 MHz, d6-DMSO): *δ* (ppm), 9.83 (s, 1H), 8.08 (d,1H), 8.01 (d, 1H), 7.63 (d, 1H), 7.51 (m, 2H), 7.41 (m, 1H), 6.97 (d, 1H), 4.15 (q, 2H), 1.40 (t, 3H). 13C NMR (100 MHz, d6-DMSO): *δ* (ppm) 167.51, 153.61, 150.22, 147.22, 134.12, 126.40, 124.88, 124.25, 122.16, 121.20, 115.93, 111.16, 63.96, 14.67. IR (KBr, cm^−1^): 3053.62(w), 2359.43(w), 2341.56(w), 1602.77(s), 1519.55(m), 1480.79(s), 1455.52(w), 1434.58(s), 1408.33(m), 1314.93(m), 1227.40(s), 1155.98(m), 1097.13(m), 967.47(s), 939.05(w), 835.99(s), 807.76(m), 755.37(s), 728.81(s), 696.45(m), 622.35(m), 550.68(w), 532.19(m), 498.99(m), 439.57(w). ESI-MS: m/z: M+: cal: 271.33, found: 272.07.

### Synthesis of cIr-Tub

4-(benzo[d]thiazol-2-yl)-2-ethoxyphenol (2.2mmol, 0.60 g) and IrCl3·3H2O (1.0 mmol, 0.35 g) were dissolved in the mixing 2-ethoxyethanol and water (3:1, v/v, 20 mL) in a flask, and refluxed for 24 h. After cooled the mixture down, the orange precipitate was filtered to give cyclometalated Ir(III) chloro-bridged dimmer. The chloro-bridged dimer (0.25 mmol, 0.34 g) and phenanthroline (1 mmol, 0.18 g) were placed in a 100 mL round bottomed flask with methanol and dichloromethane (2:1, v/v, 50 mL). The mixture was heated at 70 °C for 12 h under argon. After cooled to room temperature, NH4PF6 (2.5 mmol, 0.41 g) was added into the reaction system, and continuously stirred for 4 h. The product was purified by silica chromatography column using dichloromethane and methanol (50:1, v/v) as the eluent to obtain an orange-red solid (yield 76.5 %). 1H NMR (400 MHz, d6-DMSO): *δ* (ppm), 9.54 (s, 2H), 8.92 (d, 2H), 8.49 (d, 2H), 8.32 (s, 2H), 8.15 (q, 2H), 8.04 (d, 2H), 7.53 (s, 2H), 7.17 (t, 2H), 6.82 (t, 2H), 5.89 (s, 2H), 5.60 (d, 2H), 4.17 (m, 4H), 1.36 (t, 6H). 13C NMR (d6-DMSO, 100 MHz): *δ* (ppm), 180.8, 152.2, 151.3, 148.6, 146.8, 145.6, 144.2, 139.22, 131.4, 130.7, 130.6, 128.3, 127.5, 127.4, 125.1, 124.2, 115.7, 114.3, 110.0, 55.8, 54.5. IR (KBr, cm^-1^): 3053.63(w), 2359.43(w), 2341.56(w), 1602.77(s), 1519.55(s),1480.79(s), 1434.58(s), 1408.33(m), 1314.93(m), 1227.40(s), 1155.98(s), 1097.13(m), 967.47(s), 835.99(s), 807.76(m), 755.37(s), 728.819(s), 696.45(m), 622.35(s), 550.68(m), 522.19(m), 498.99(m); ES-MS: cal:913.13, found: 913.33. cIr-1 and cIr-2 were obtained by the same synthetic procedures used for cIr-Tub. cIr-1, 1H NMR (400MHz, d6-DMSO6): *δ* (ppm), 8.95 (d, 2H), 8.46 (d, 2H), 8.36 (s, 2H), 8.16 (dd, 2H), 8.10 (d, 2H), 7.58 (s, 2H), 7.24 (t, 2H), 6.92 (t, 2H), 5.80 (s, 2H), 5.75 (d, 1H), 3.85 (s, 6H), 3.14 (s, 6H). 13C NMR (d6-DMSO, 100 MHz): δ (ppm), 180.84, 152.19, 151.34, 148.6, 146.87, 145.64, 144.19, 139.22, 131.44, 130.69, 130.61, 128.31, 127.50, 127.37, 125.08, 124.23, 115.77, 114.26, 110.04, 55.87, 54.49. IR (KBr, cm^−1^): 3081.15(w), 3053.77(w), 2965.48(w), 2939.19(w), 2841.95(w), 2360.93(w), 1599.53(s), 1522.07(s), 1482.18(s), 1431.81(s), 1420.32(s), 1262.25(s), 1243.37(m), 1146.58(s), 1018.69(s), 875.53(m), 808.05(s), 764.67(s), 733.48(m), 696.89(s), 652.47(m), 563.84(w); ES-MS: cal. 913.15, found 913.33. cIr-2, 1H NMR (400MHz, d6-DMSO6): *δ* (ppm), 8.94 (d, 2H), 8.52 (d, 2H), 8.36 (s, 2H), 8.19 (dd, 2H), 7.97 (d, 2H), 7.75 (d, 2H), 7.13 (t, 2H), 6.84 (dd, 2H), 6.53 (d, 2H), 5.64 (d, 2H), 5.55 (d, 2H), 4.58 (s, 2H), 3.13 (s, 4H), 3.01 (dd, 4H), 2.63 (s, 6H). 13C NMR (d6-DMSO, 100 MHz): δ (ppm), 179.75, 152.44, 151.18, 150.77, 149.03, 146.94, 138.99, 130.57, 130.06, 128.26, 128.17, 127.30, 127.15, 127.09, 124.14, 123.76, 115.23, 113.58, 106.92, 58.04, 53.89. IR (KBr, cm^−1^): 3429(w), 3069(w), 2919(w), 2878(w), 2359(w), 2346(w), 1580(s), 1492(m), 1456(s), 1432(s), 1396(s), 1367(w), 1347(w), 1321(w), 1261(m), 1246(m), 1201(m), 1153(s), 1043(s), 991(m), 837(s), 781(w), 755(s), 723(s), 723(s), 669(w), 645(w), 600(w), 557(s), 446(w), 428(w); ESI-MS: cal. 939.43, found 939.16.

### Cell culture

HepG2 (hepatocellular liver carcinoma) cells purchased from Shanghai Bioleaf BioBiotech. Co. Ltd. The cells were incubated in Dulbecco’s Modified Eagle’s medium (DMEM) containing 10 % FBS and 1% antibiotics (penicillin and streptomycin), maintained at 37 °C in an atmosphere of 5 % CO2 and 95 % air.

### Cytotoxicity assays

The cytotoxicity of Ir-Tub towards HepG2 cells was determined by 5-dimethylthiazol-2-yl-2,5-diphenyltetrazolium bromide (MTT) assay. The exponentially grown HepG2 cells were seeded in triplicate into 96-well plates at 104 cells per well. After 48 h, the cells were treated with Ir-Tub at different concentrations (1 *μ*M, 5 *μ*M, 10 *μ*M, and 20 *μ*M) and incubated for 24 h. After that time, the media was removed, and the cells were rinsed once with PBS and placed with fresh media. Subsequently, cells were treated with 5 mg/mL MTT (10 μL/well) and incubated for an additional 4 h (37 oC, 5 % CO2). After MTT medium removal, the formazan crystals were dissolved in DMSO (100 μL/well). The plate was incubated for 10 min while shaking it with an oscillator. The absorbance was measured at 450 nm using a microplate reader (SpectraMax Paradigm).

### Tissue sectioning

Fresh mouse organs were put in liquid nitrogen surrounding by isopentane. The slices (20 μm thickness) were obtained by Leica CM3050S freezing microtome. Before imaging, the slices were stained with cIr-Tub (10 μM) for 60 min at 37 °C, followed by washing with PBS buffer three times. Note: All procedures involving animals were approved by and conformed to the guidelines of the Southwest University Animal Care Committee, College of Pharmaceutical Sciences. We have taken great efforts to reduce the number of animals used in these studies and also taken the effort to reduce animal suffering from pain and discomfort.

### Immunofluorescence

Simply, the fixed organ slice or cells was exposed to 0.5 %Triton X-100 for 5 min and washed by PBS buffer three times. After incubated with 100 mM glycine for 15 min at room temperature, the slice was covered with 1 % BSA for 1 hour in order to close non-specific binding sites. Subsequently, the primary antibody was added onto the slice, and kept in the refrigerator of 4°C for 12 h. Finally, the primary antibody-binding slices were incubated with fluorescent second antibodies. The imaging of slices was carried out after washed by PBS buffer three times.

### STED super-resolution image

The compound was excited under STED laser (doughnut laser: 595nm), the emission signals were collected using HyD reflected light detectors (RLDs) with 2048*2048 pixel and *100 scanning speed. The STED micrographs were further processed ‘deconvolution wizard’ function using Huygens Professional software (version: 16.05) under authorized license. The area radiuses were estimated under 0.02 micros with the exclusion of 100 absolute background values. Maximum iterations were 40-time, signal-to-noise ration 20 was applied, with quality threshold 0.05; iteration mode: Optimized; Brick layout: Auto.

### Correlative light-electron microscopy (CLEM)

The ultra-thin (100-120nm) sections from the above process were moved to ZEISS LSM800 with Airyscan to obtain light microscope data. Electron microscopic imaging was taken under GeminiSEM 500 scanning transmission electron microscopy. To correlative, ZEISS ZEN connect module was chosen to find the same scan region under LM and EM, and ZEISS ZEN shuttle and find module was used merge two images together, the process was described as follow: (1) Set up a connect project in ZEN Blue, and take an overview bright field image of the whole copper grid under LSM800 with the objective of EC Epiplan 5X/0.13. (2) Choose different regions of interest to scan high magnification fluorescent image in the connected project that first set up. (3) Transfer the copper grid sample to ZEISS GeminiSEM 500 and use the ZEN connect module to open the project data saved in LSM800. (4) Take a large field of view image on GeminiSEM 500 and align those two-overview images. (5) Click the position where the fluorescent image takes; the stage will automatically move to the position where the FL image is. (6) Take the EM image for the sample, and merge the two images by using the ZEISS ZEN shuttle and find module based on the 3-points method.

**Figure S1:**
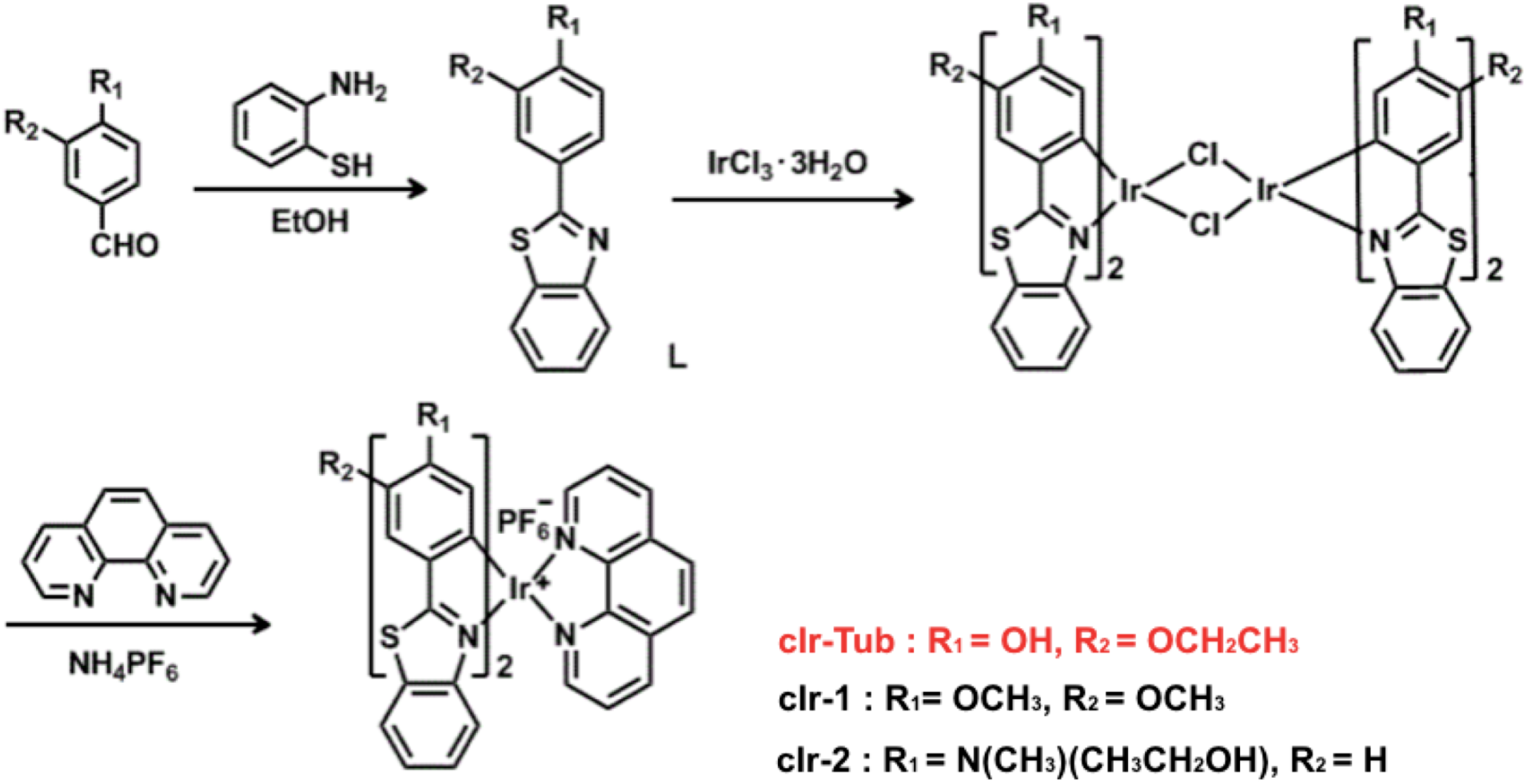
The synthetic routes of cIr-Tub and its analogous compounds (cIr-1, cIr-2).

### Chemical analysis by Energy-Filtered TEM

The chemical analysis investigation of the specimen was performed using a JEOL JEM-2200FS TEM equipped with a field emission gun (FEG) at 200 kV, and an in-column energy Omega filter. Images were taken at a collection angle of 24.282 mrad. The microscope was used in energy-filtered transmission electron microscopy (EF-TEM) mode in order to perform elemental analysis, and, thus, fine structure imaging. The software used for image acquisition and processing was the Digital Micrograph(tm) software (version 3.20). Images were recorded using a charge-coupled device (CCD) camera US1000XP from Gatan. The Iridium present in the cells was identified by exciting the atomic O shell (50 eV) with a slit width of 6.0 eV. The 3-window technique was employed. This technique entails two images before the ionization edge and one after. The pre-edge images are used to compute the approximate background contained in the post-edge window. Once the background is determined and removed, the subsequent map displays a signal that is proportional to the element concentration in the sample. The first pre-edge second pre-edge and post-edge energies were 39 eV, 45 eV, and 53 eV respectively. The unfiltered TEM image and the false colour chemical mapping were superimposed using ImageJ software.

**Figure S2:**
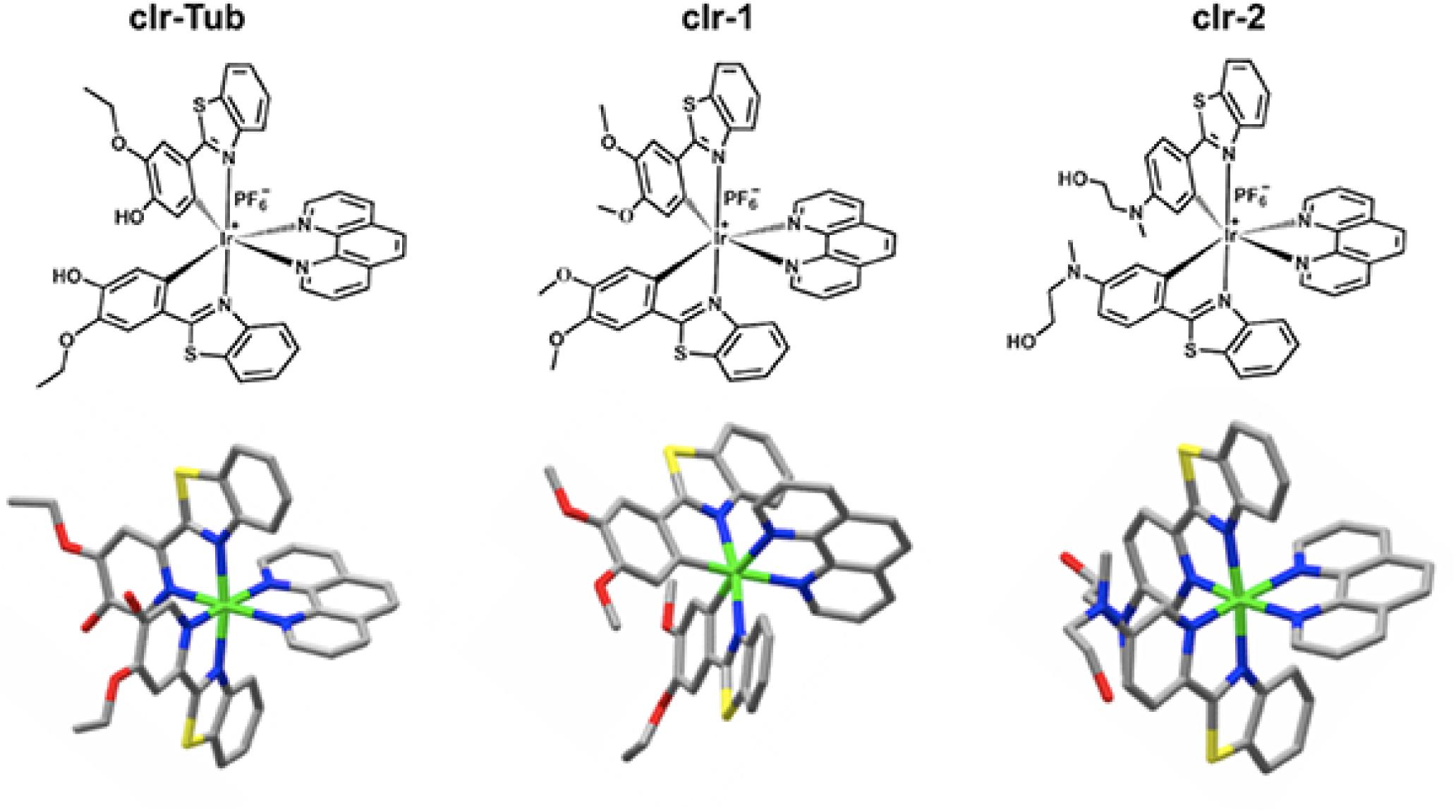
Chemical and stereo structures of cIr-Tub, cIr-1 and cIr-2 complexes.

**Figure S3:**
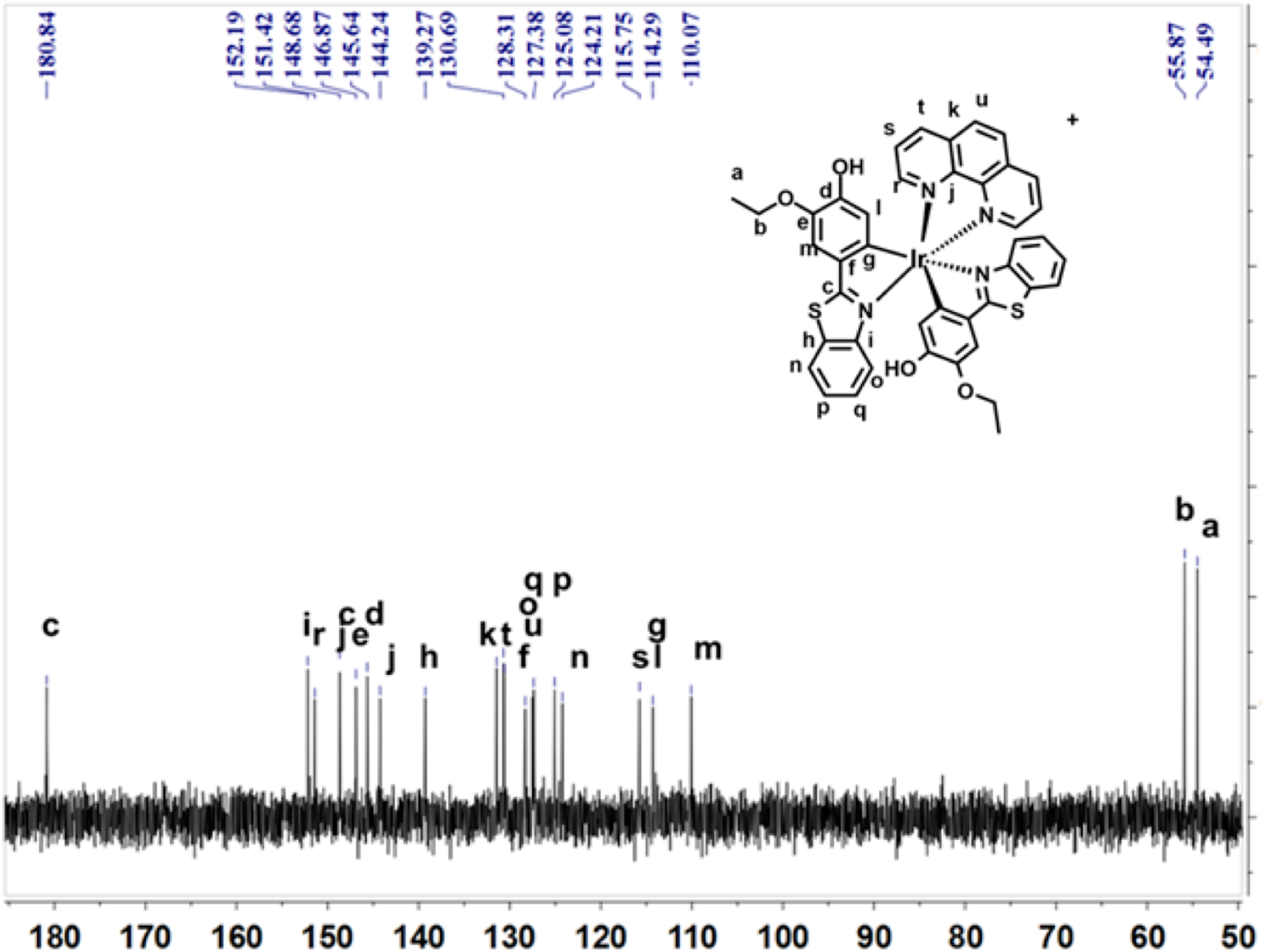
^13^C NMR spectra of cIr-Tub.

**Figure S4:**
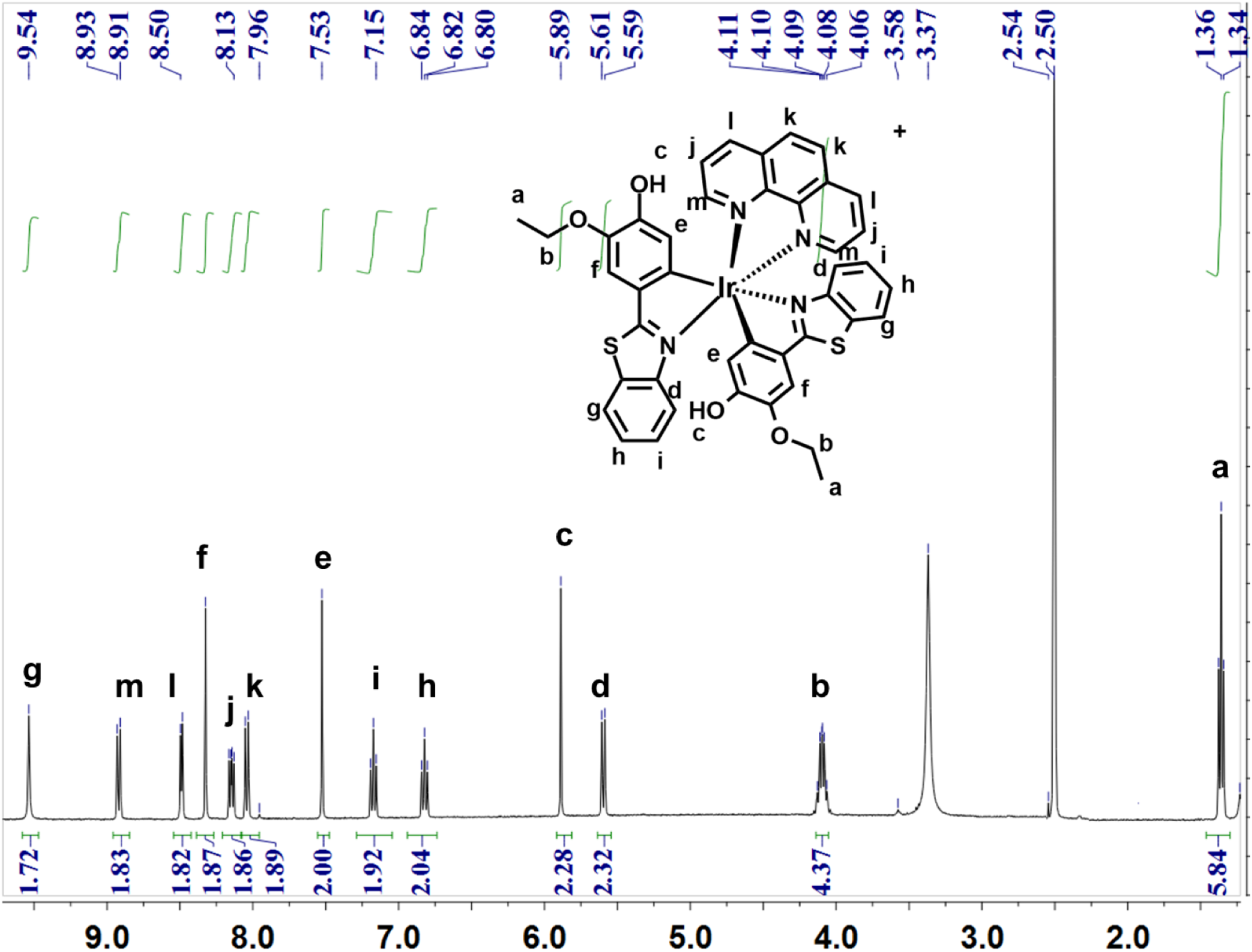
^1^H NMR spectra of cIr-Tub.

**Figure S5:**
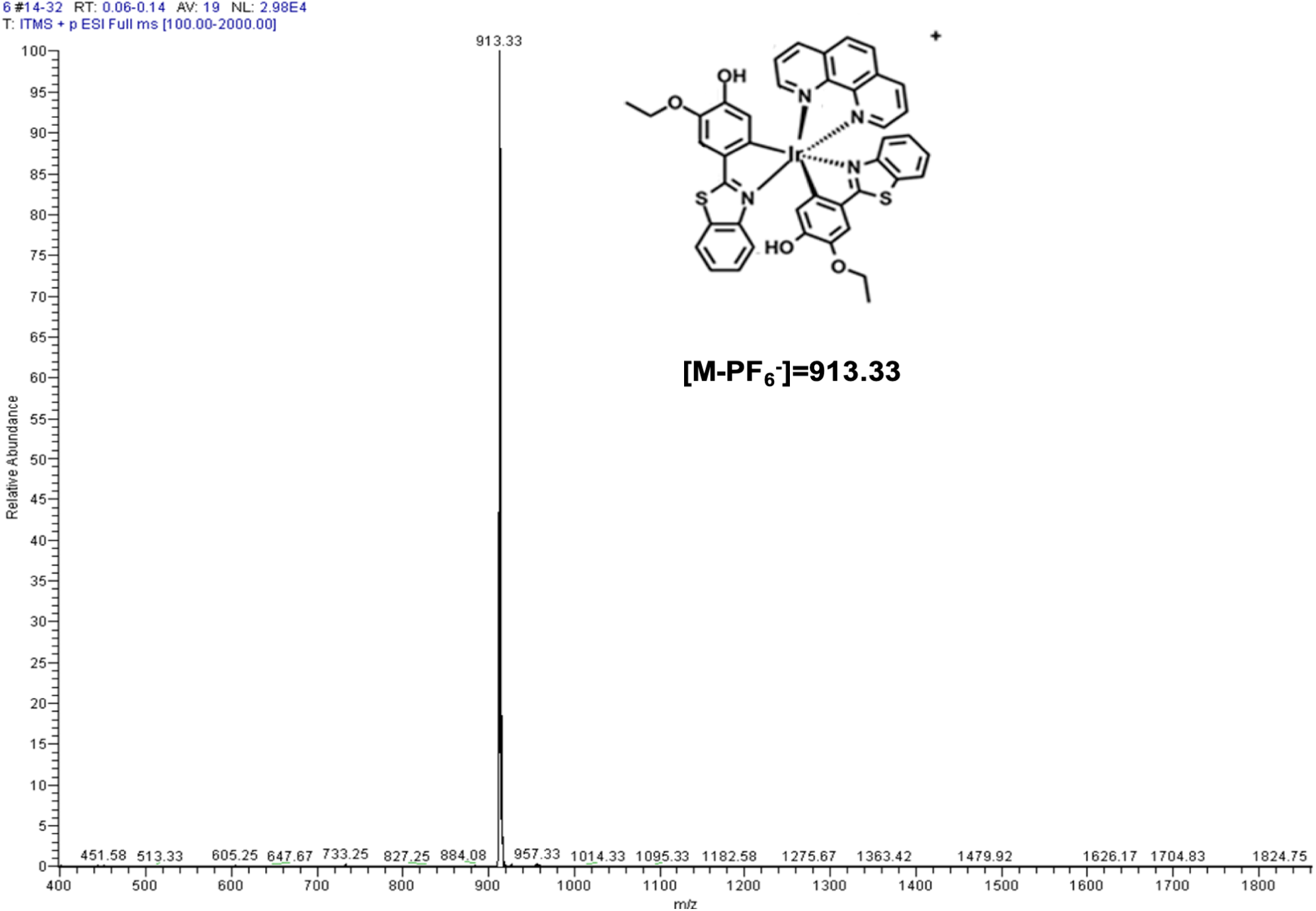
Mass spectrum of cIr-Tub.

**Table S1:**
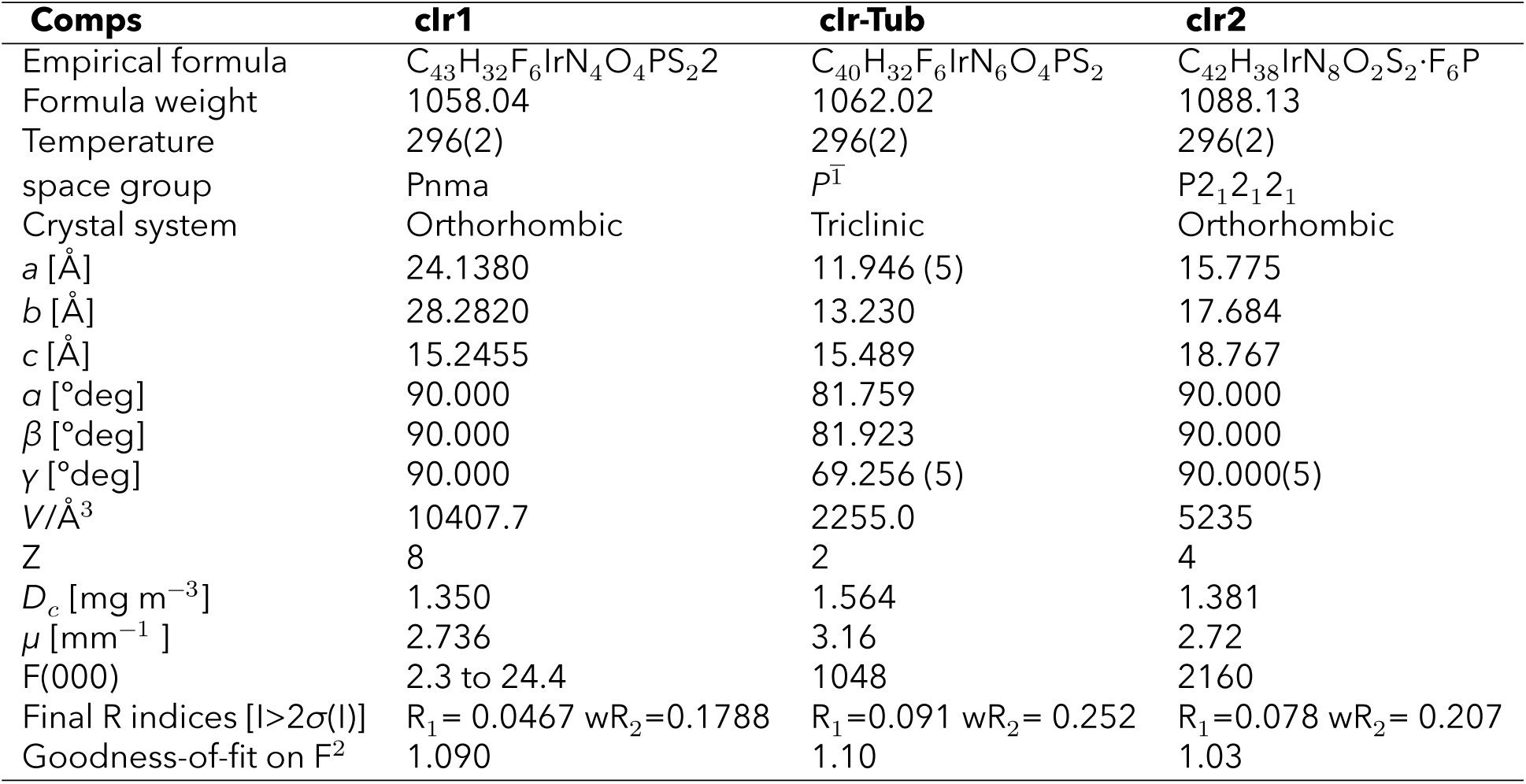
Crystal data collection and structure refinement of cIr1, cIr-Tub and cIr2.

**Table S2:** Photophysical properties of cIr-Tub in different solvents and bound to tubulin.

| | Absorption peak<br>$\lambda_{max}^{exc}$ [nm] | $\epsilon_{max}$<br>$10^4$ [mol<sup>-1</sup> L cm<sup>-1</sup>] | Emission peak<br>$\lambda_{max}^{em}$ [nm] | Stoke shift<br>$\Delta\nu_{max}^0$ [nm] | Quantum yield<br>$\phi$ | Fluorescence life time<br>$\tau^f$ [ns] |
| --- | --- | --- | --- | --- | --- | --- |
| Dichloromethane | 437 | 1.42 | 593 | 156 | 0.005 | 20.46 |
| Tetrahydrofuran | 440 | 1.58 | 611 | 171 | 0.002 | 9.88 |
| Ethanol | 431 | 1.23 | 609 | 178 | 0.001 | 9.65 |
| PBS | 443 | 1.77 | 560 | 117 | <0.001 | 9.77 |
| Ir-Tub+tubulin | 445 | 1.88 | 563 | 118 | 0.044 | 18.33 |

**Figure S6:**
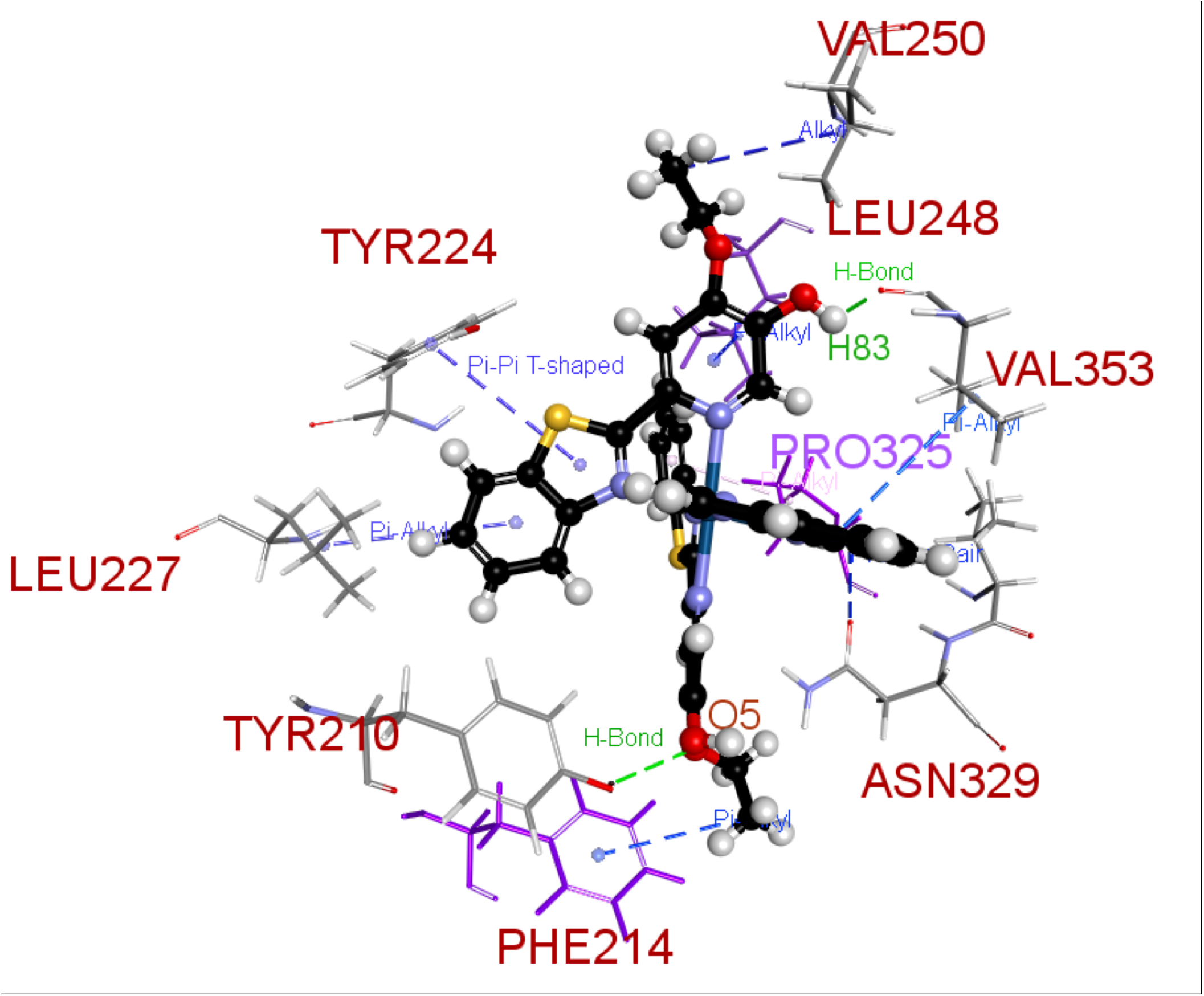
Detailed interaction between cIr-Tub complex with tubulin protein (Phe214, Asn329, Pro325, Val353, Leu248, Val250, Tyr224, Leu227 and Tyr210) by molecular docking simulation.

**Table S3:** Binding energy of cIr-Tub compare with vinblastine that interaction with tubulin protein.

| Ligand Name | Binding Energy | Ligand Energy | Tubulin Energy<br>[Kcal mol<sup>-1</sup>] | Complex Energy | Entropy |
| --- | --- | --- | --- | --- | --- |
| Vinblastine | -7.6413 | 234.0482 | -85087.7320 | -84861.3251 | 21.8125 |
| clr-Tub | -199.9281 | 1596.4059 | -85087.7320 | -83691.2542 | 21.9300 |

**Figure S7:**
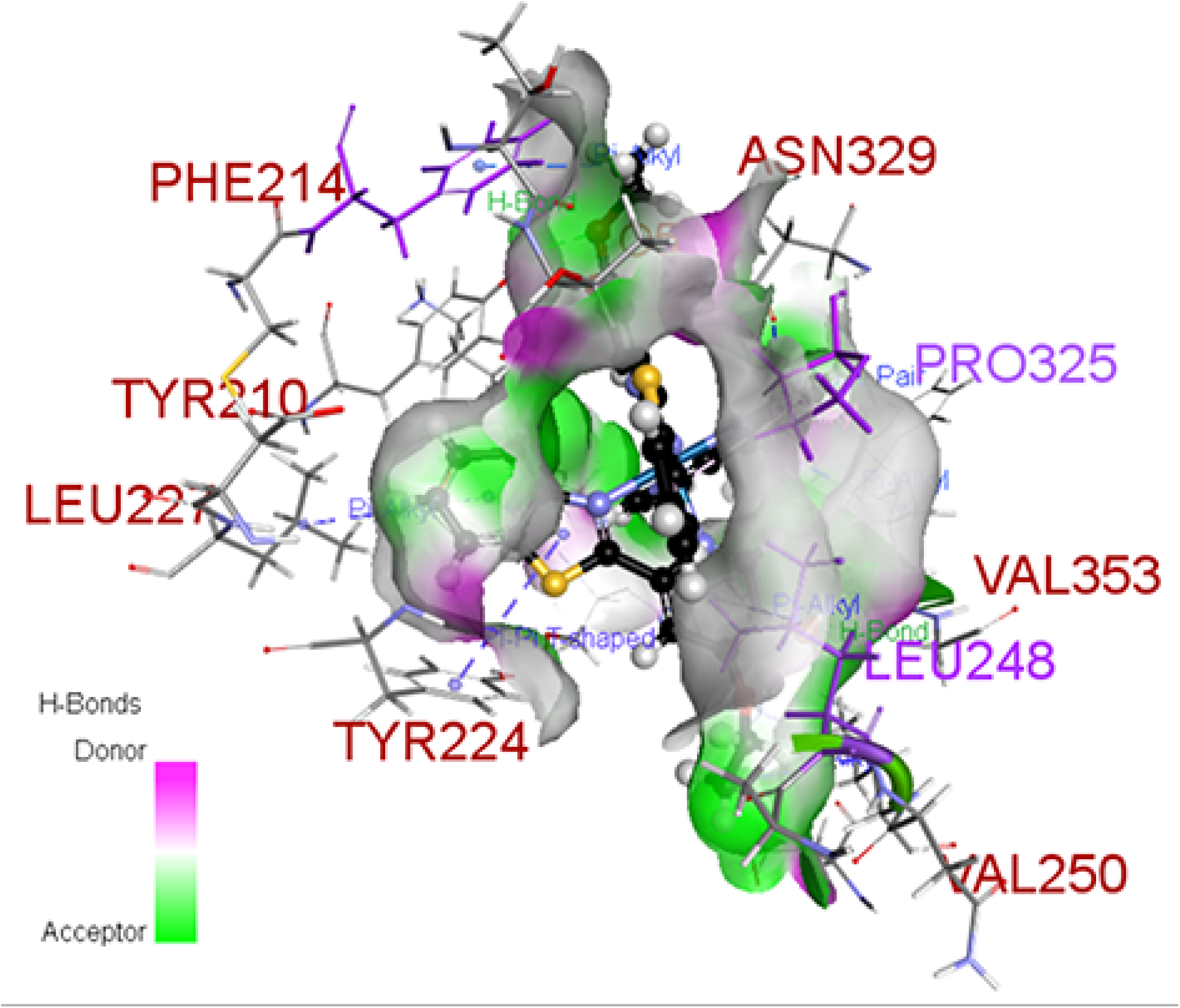
Computer modelling of Hydrogen bonds map between cIr-Tub complex and tubulin.

**Figure S8:**
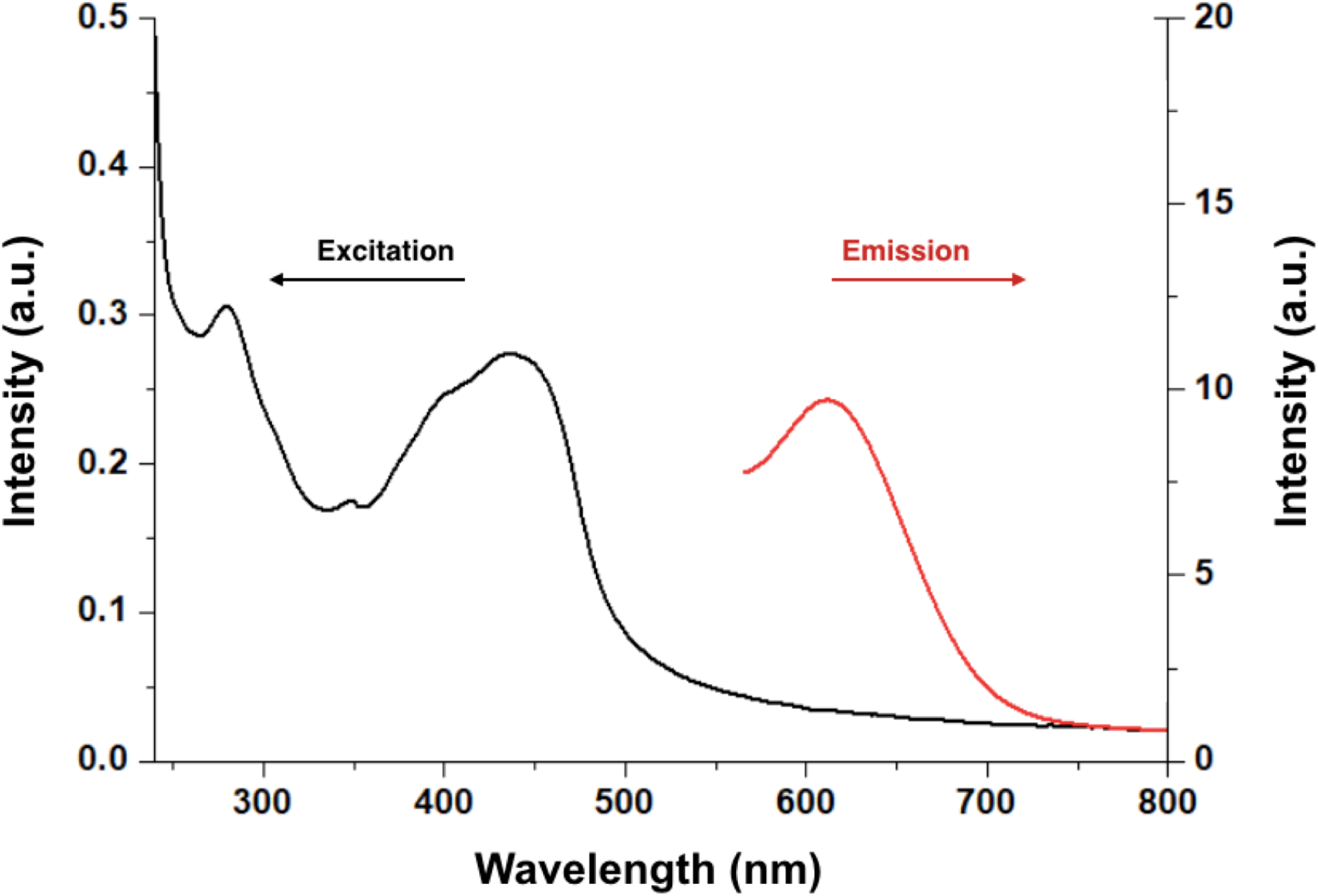
UV-vis absorption spectra (black) and fluorescence spectra (red) of Ir-Tub in PBS (excited at 445 nm).

**Figure S9:**
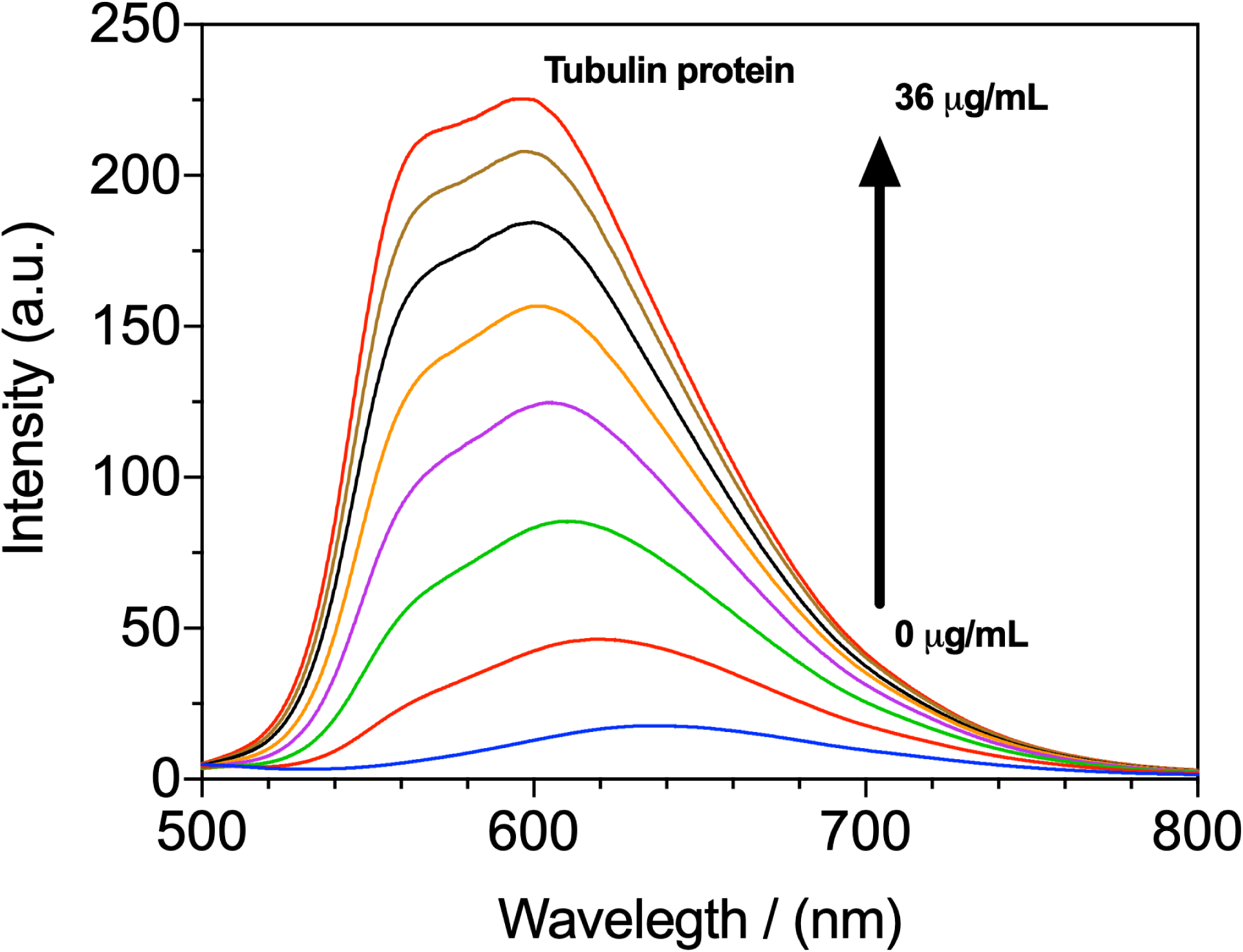
Fluorescent spectra of cIr-Tub complex upon addition of tubulin protein. Excitation wavelength at 450 nm.

**Figure S10:**
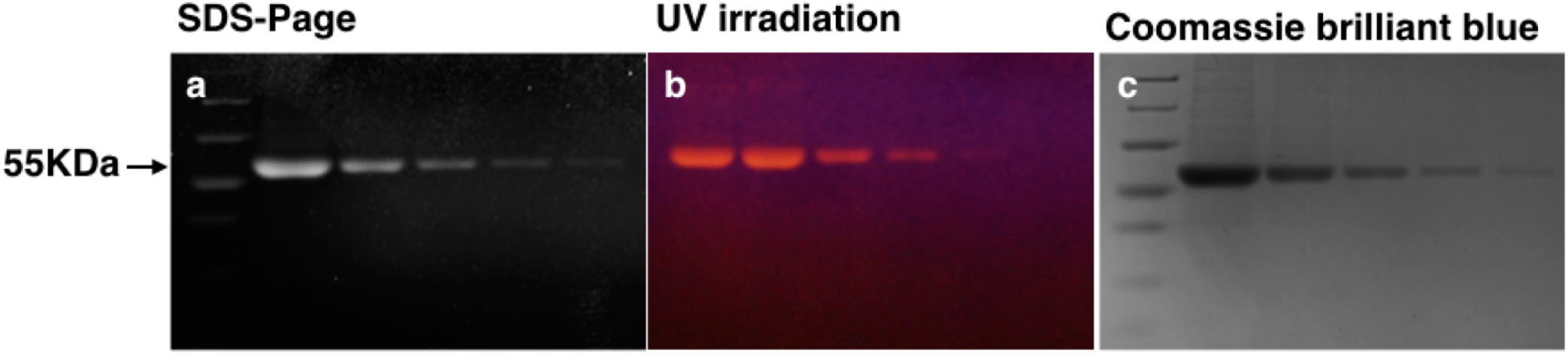
(**a**) The SDS-PAGE gel showing tubulin-binding by pre-incubated with cIr-Tub complex with decreased concentration of purified tubulin. (**b**) same gel exposed under UV light also (**c**) incubated with coomassie brilliant blue (right).

**Figure S11:**
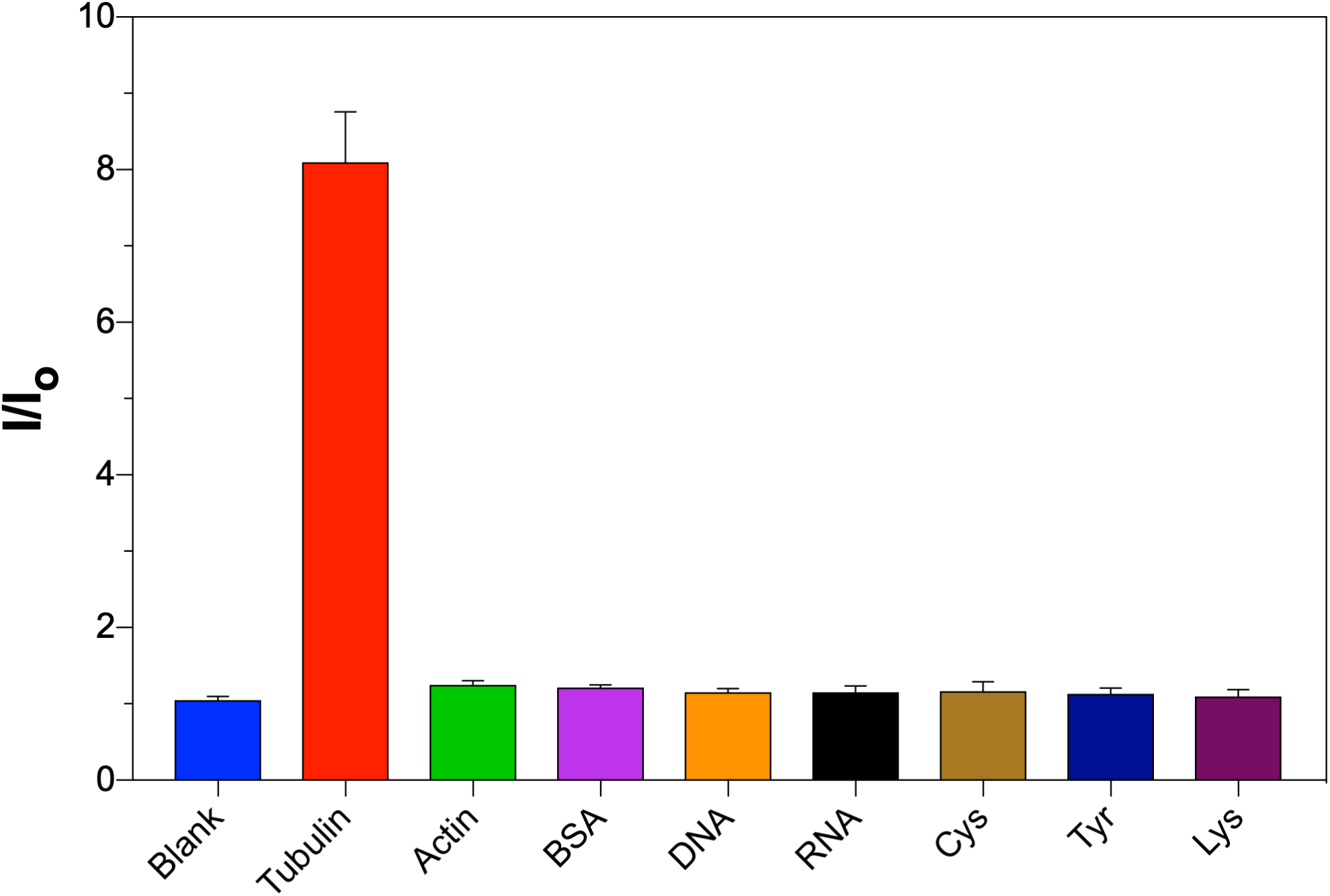
The fluorescent selectivity of Ir-Tub (10 *μ*M) to various biomolecules: Blank, 30 μg/mL tubulin, 30 *μ*g/mL actin, 30 *μ*g/mL bovine albumin, 10 *μ*M dsDNA, 10 *μ*g/mL ssRNA and amino acids (Cys, Tyr, Lys, 200*μ*M). The fluorescent intensity was measured by the excitation with 450 nm and the collection at 600 nm. The bars represent the mean errors from three individual experiments.

**Figure S12:**
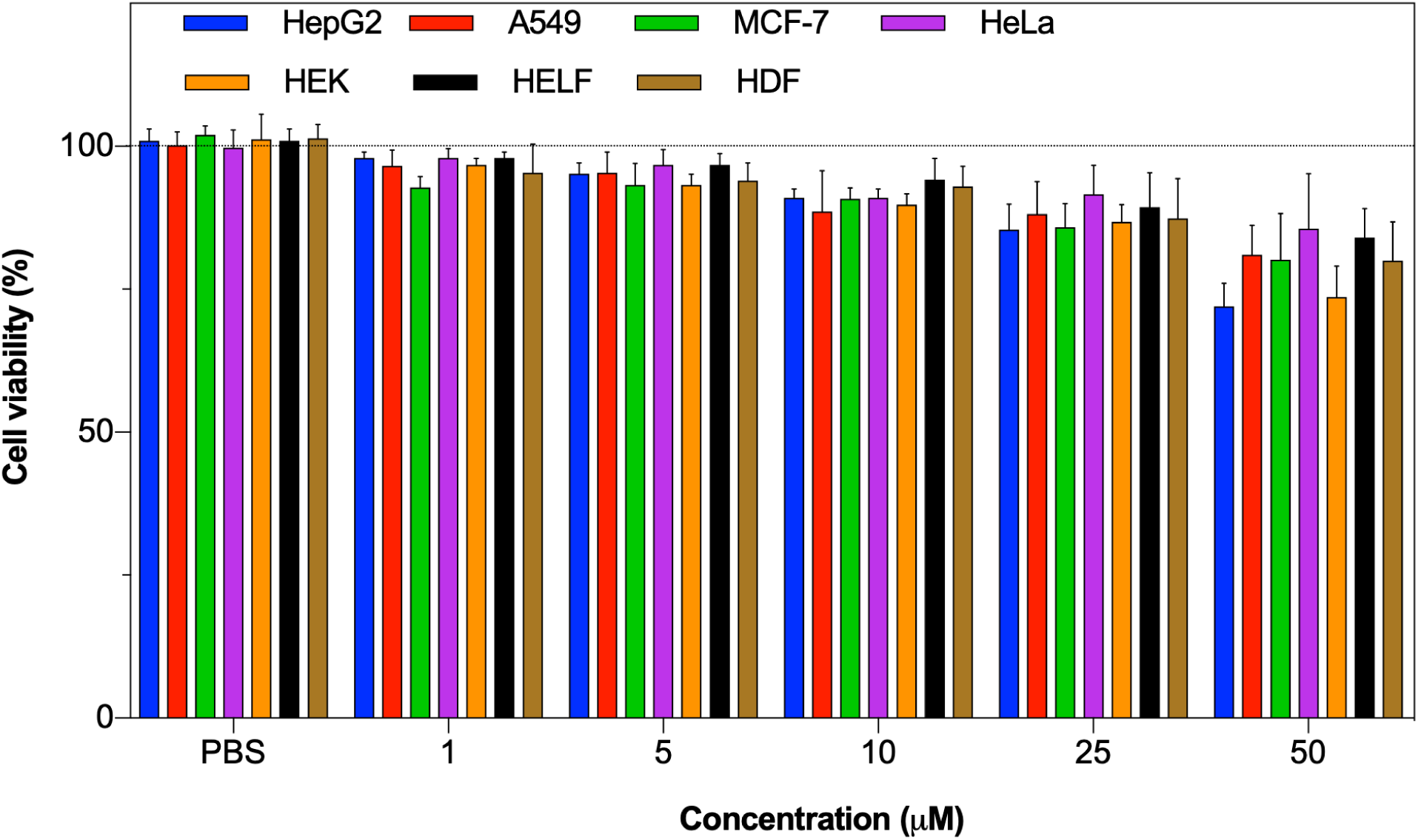
MTT assay using cIr-Tub complex against cancer cells including HepG2, A549, MCF-7 and HeLa and normal cells including HEK, HELF and HDF, incubation time is 12 hours.

**Figure S13:**
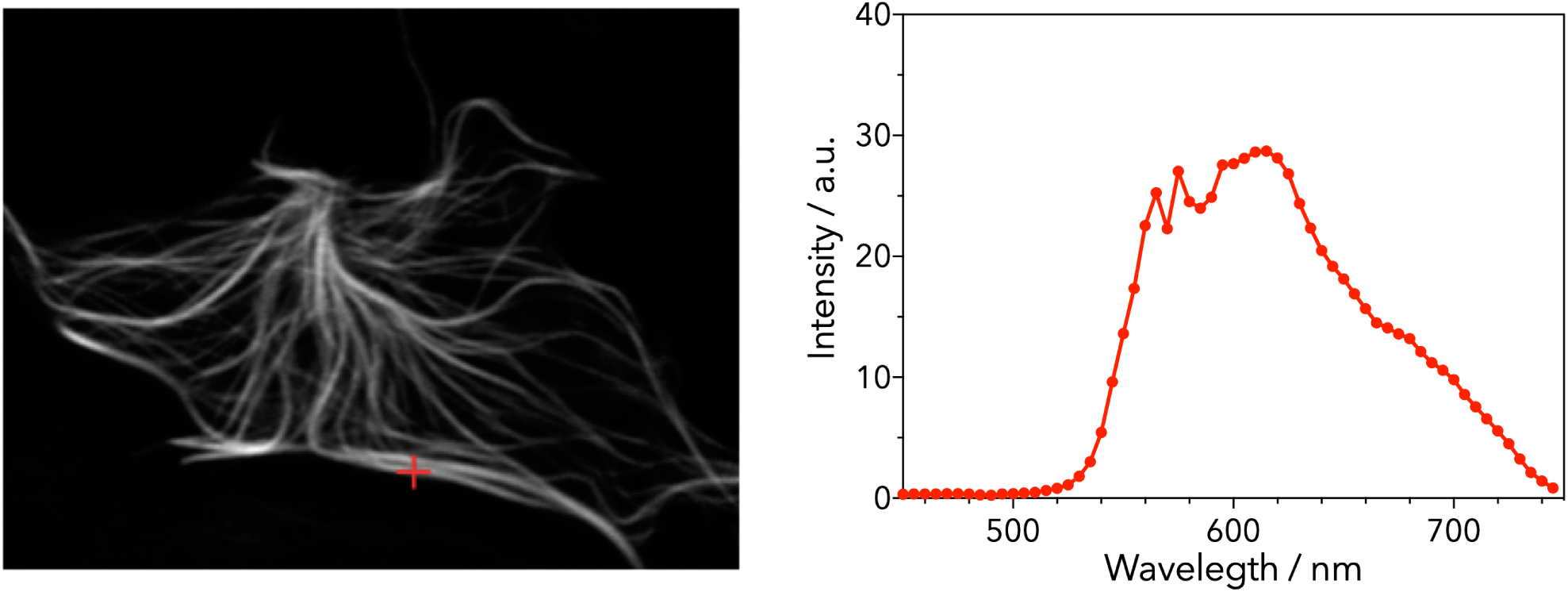
Lambda-scanning (*λ*-scan) experiment using cIr-Tub complex incubated live cells measure the emission intensity across a range of wavelengths (450nm-750nm, interval=5nm).

**Figure S14:**
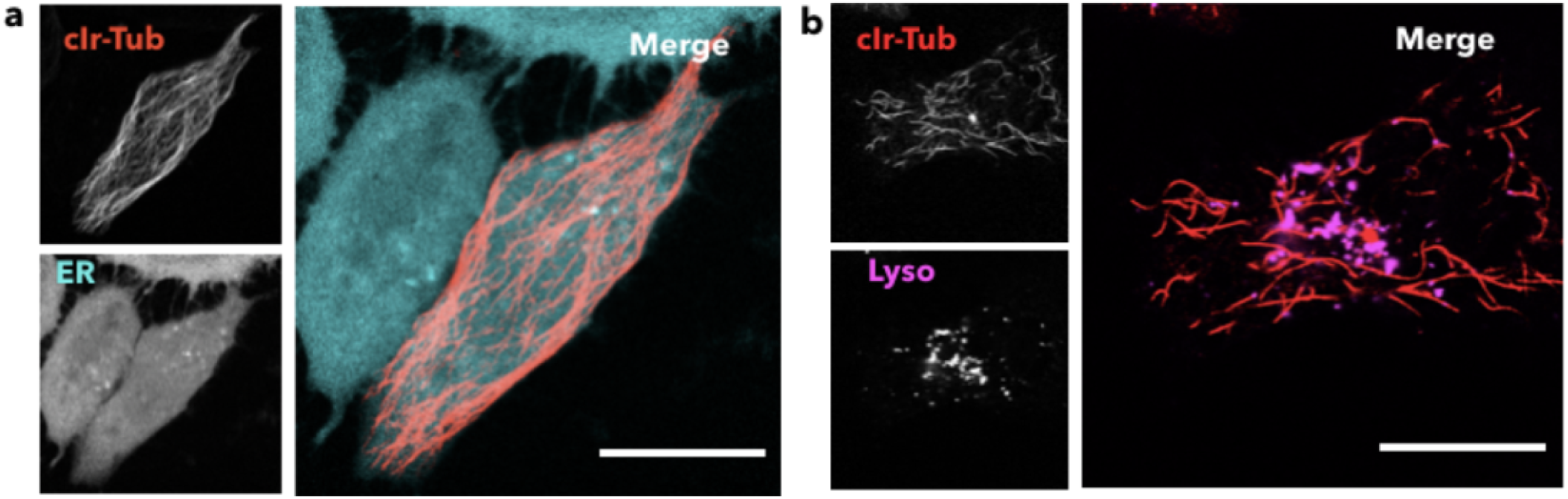
Live cell co-localization of cIr-Tub with (**a**) endoplasmic reticulum and (**b**) lysosomes stained with ER tracker and lysotracker to stain respectively. Scale Bar = 10 *μ*m.

**Figure S15:**
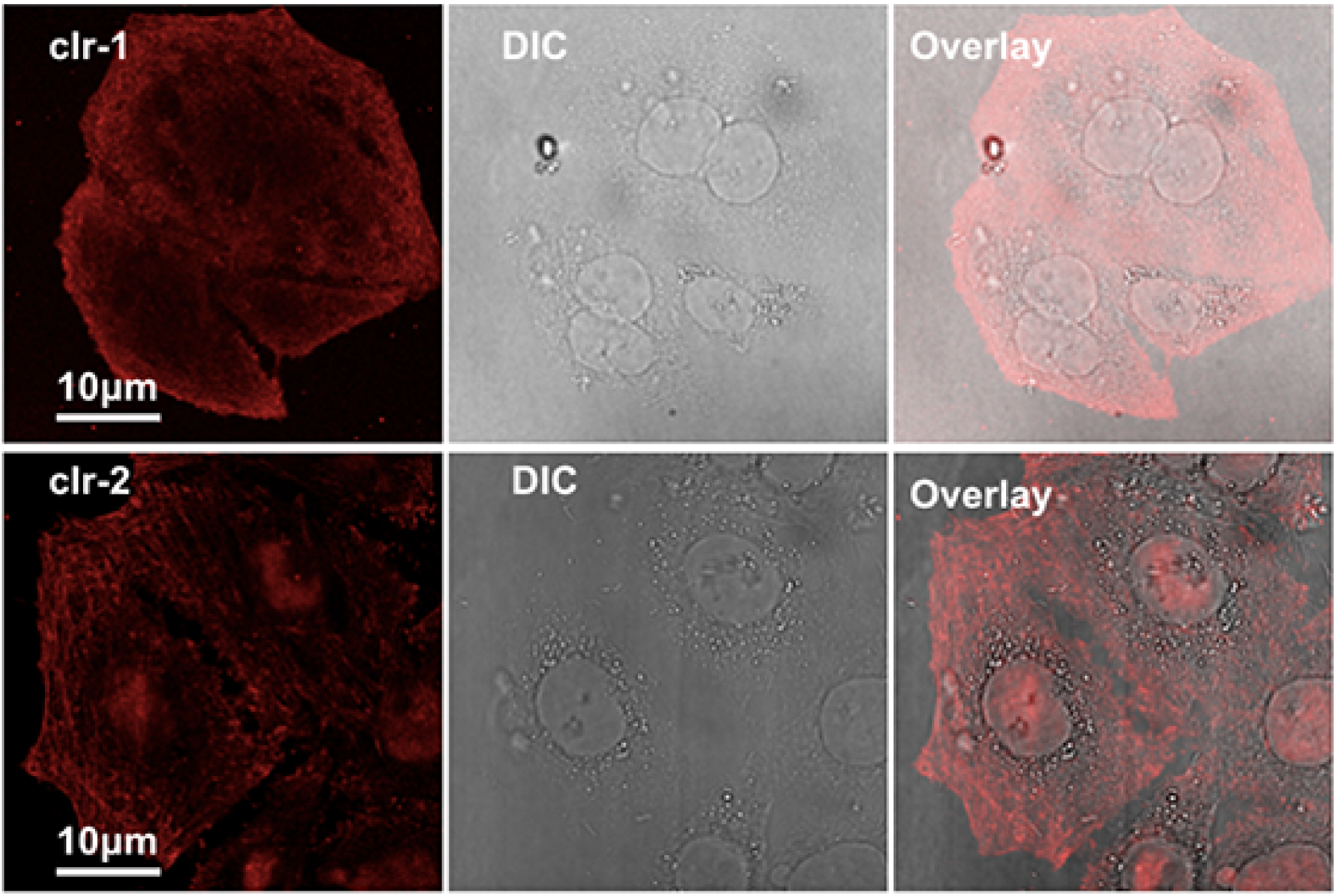
HepG2 cell lines incubated with cIr-1 and cIr-2 complex (5 μM), then directly imaged under confocal microscopy.

**Figure S16:**
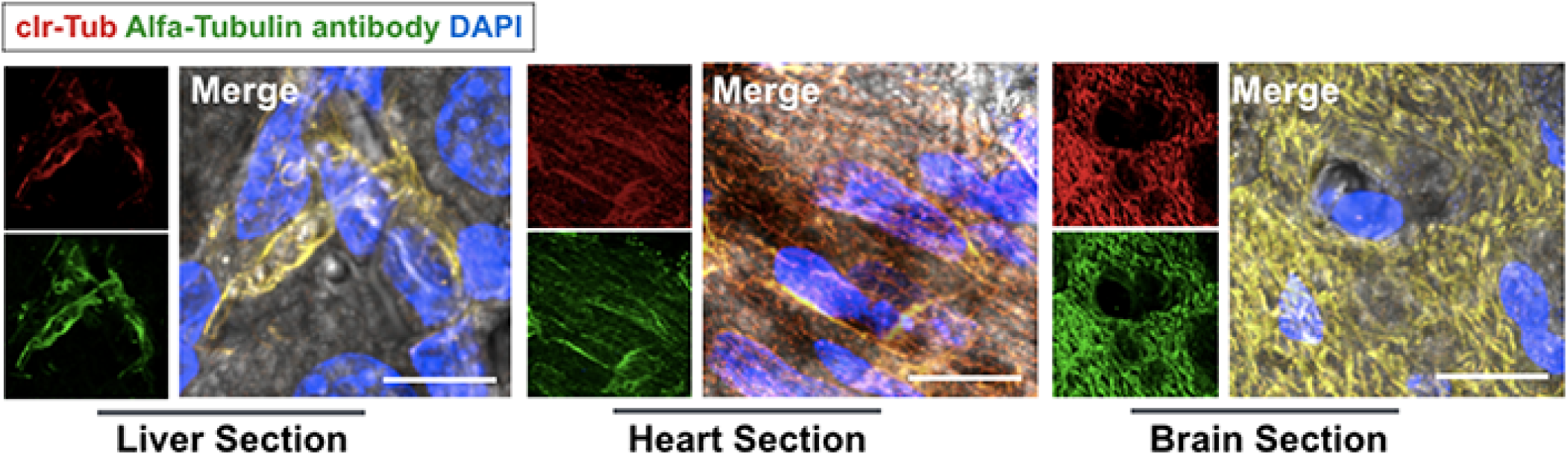
Tissue slices sectioned from liver, heart, and brain were incubated with cIr-Tub complex for 30min and treated for immunofluorescence using alfa-tubulin antibody, the nuclear were highlighted using DAPI.

**Figure S17:**
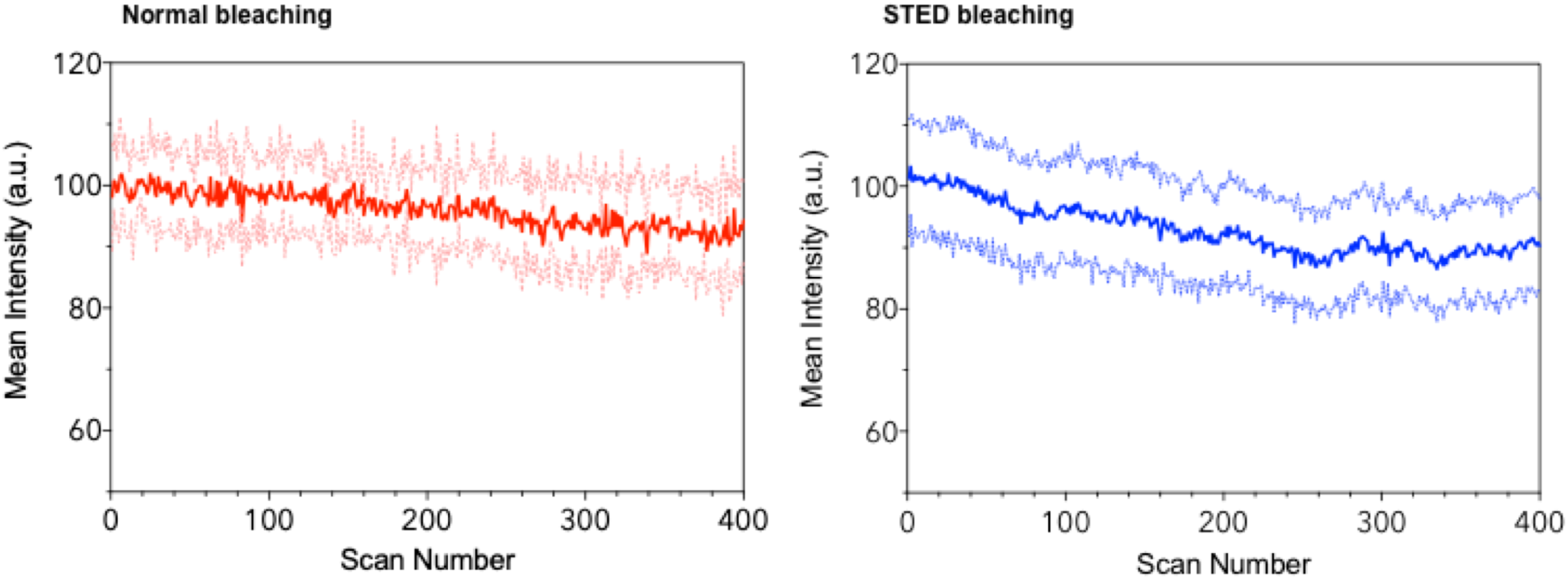
Photobleaching experiments using cIr-Tub HepG2 treated cells against normal (left) and STED (right) scanning. Normal scanning Laser: 405nm (16 mW); STED scanning laser: 405 nm (16 mW) + 595nm donut laser (6 mW).

**Figure S18:**
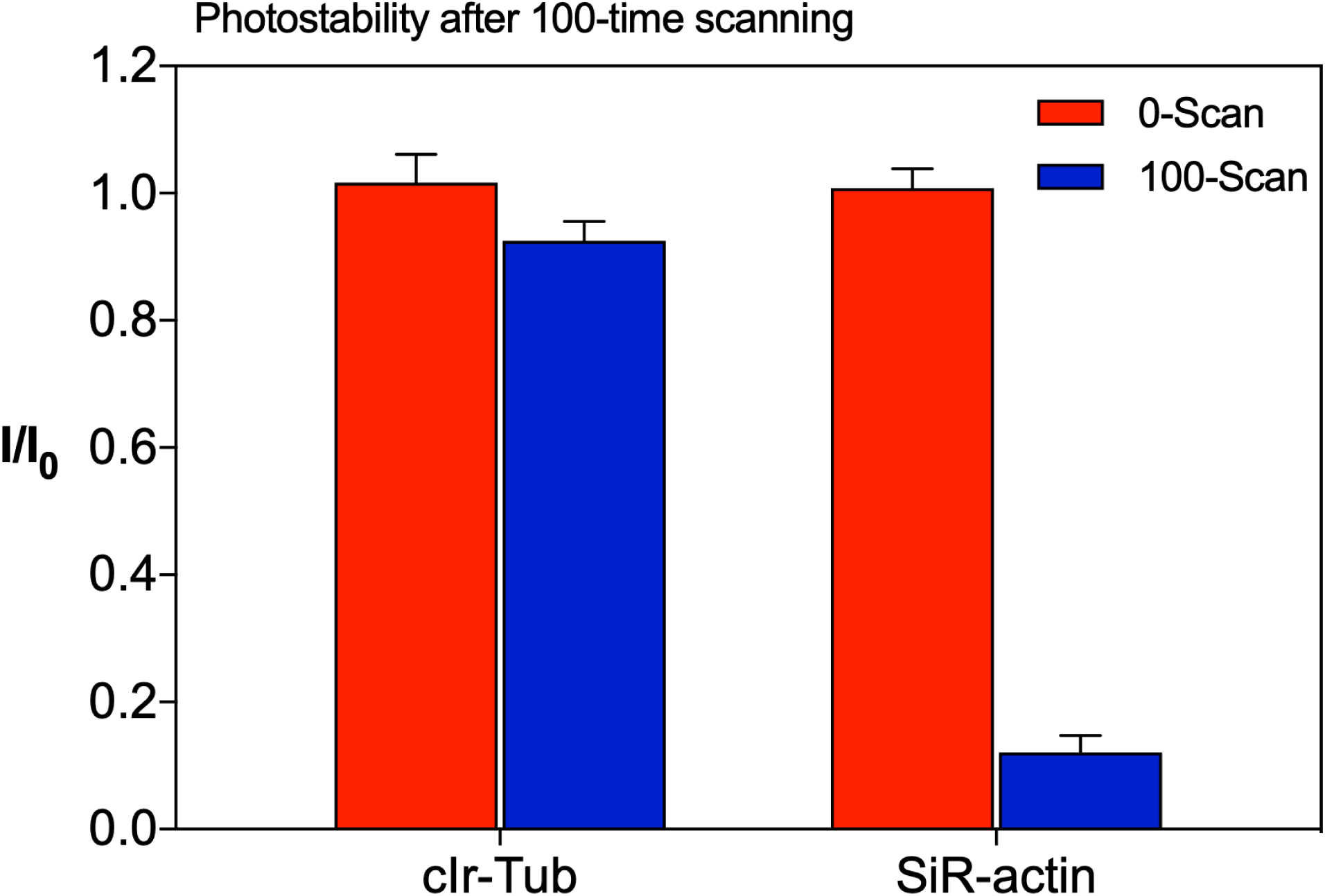
Photobleaching analysis using cIr-Tub and SiR-Tubulin treated HepG2 cells after 100 times continued STED scanning. STED laser for cIr-Tub: 405nm(16 mW) + 595nm donut laser (6 mW); STED laser for SiR-Tubulin: 640 nm(16 mW) + 660 nm donut laser (6 mW).

**Figure S19:**
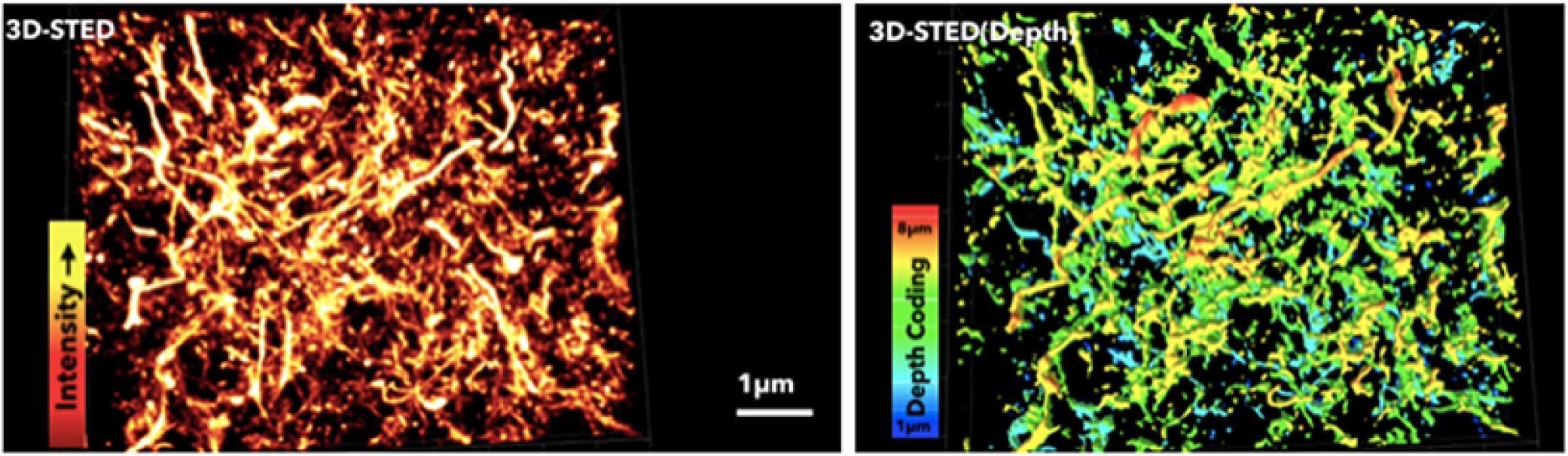
3D-STED and 3D-STED depth images from mouse brain hippocampus region, showing complex neuronal microtubule network.

**Figure S20:**
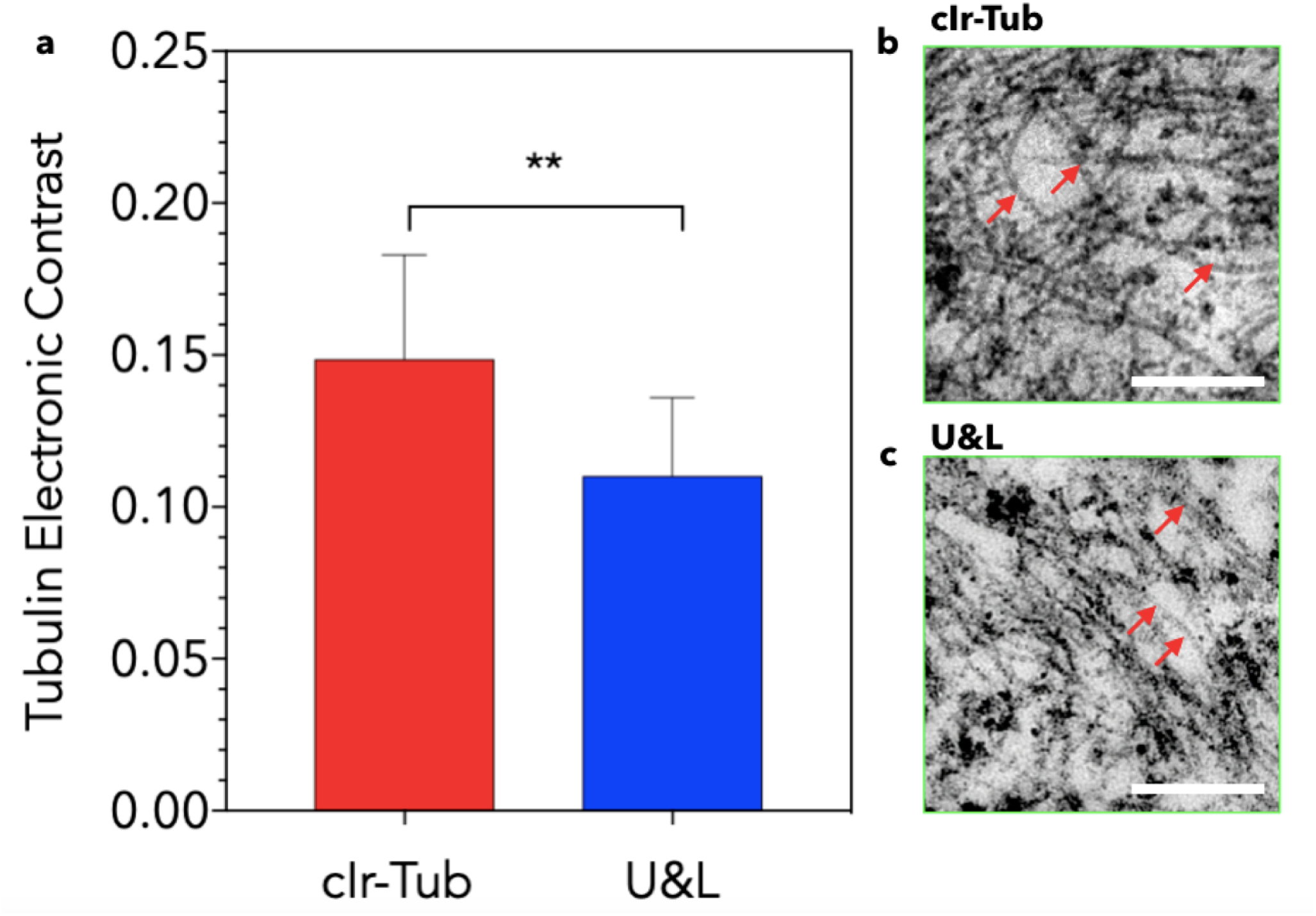
Electron microscopic contrast analysis using Michelson’s equation 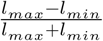 and their correspondence micrographs using **(b)** cIr-Tub complex and **(a)** classic uranyl acetate followed by staining with Reynold’s lead citrate (U&L). Scale Bar = 2 *μ*m. Red arrows indicate microtubules in cells.

